# Melanocortin-responsive Kiss1 neurons of the arcuate nucleus drive energy expenditure through glutamatergic signaling to the dorsomedial hypothalamus

**DOI:** 10.1101/2025.06.25.661567

**Authors:** Rajae Talbi, Todd L. Stincic, Nicole Lynch, Encarnacion Torres, Kaitlin Ferrari, Choi Ji Hae, Elizabeth Medve, Karol Walec, Samuel T. Zdon, Sidney A. Pereira, Carrie E. Mahoney, Martha A. Bosch, Larry Zweifel, Oline K. Rønnekleiv, Natalia L.S. Machado, Martin J. Kelly, Víctor M. Navarro

**Affiliations:** Harvard Medical School, Boston, MA, USA; Division of Endocrinology, Diabetes and Hypertension, Department of Medicine, Brigham and Women’s Hospital, Boston, MA, USA; Department of Chemical Physiology and Biochemistry, Oregon Health & Science University, Portland, OR, USA; Division of Neuroscience, Oregon National Primate Research Center, Beaverton, OR, USA; Harvard Program in Neuroscience, Boston, MA, USA; Department of Biology, Appalachian State University, Boone, USA; Department of Neurology, Beth Israel-Deaconess Medical Center, Boston, MA, USA; Department of Pharmacology and Physiology, Faculty of Medicine, Université de Montréal, Montréal, QC, Canada; Centre d’Innovation Biomédicale (CIB), Faculty of Medicine, Université de Montréal, Montreal, QC, Canada; Department of Psychiatry and Behavioral Sciences, University of Washington, Seattle, WA 98195, USA; Department of Pharmacology, University of Washington, Seattle, WA 98195, USA

**Author notes:** Corresponding authors. Victor M. Navarro, PhD, Division of Endocrinology, Diabetes and Hypertension, Department of Medicine, Brigham and Women’s Hospital and Harvard Medical School, 221 Longwood Avenue, Room 219, Boston, Massachusetts 02115. Martin J. Kelly, PhD, Department of Chemical Physiology and Biochemistry, Oregon Health & Science University, 3181 SW Sam Jackson Park Road, Portland, OR 97239.

**Keywords:** Kiss1 neurons, melanocortin system, POMC neurons, MC4R, energy expenditure, DMH, Vglut2, metabolism

## Abstract

Energy expenditure (EE) is essential for metabolic homeostasis, yet its central regulation remains poorly understood. Here, we identify arcuate Kiss1 neurons as key regulators of EE in male mice. Ablation of these neurons induced obesity, while their chemogenetic activation increased brown adipose tissue (BAT) thermogenesis without affecting food intake. This action is mediated by glutamatergic projections from Kiss1^ARC^ neurons to CART/Lepr-expressing neurons in the dorsomedial hypothalamus, which activate the raphe pallidus-BAT pathway. CRISPR-mediated deletion of the vesicular glutamate transporter 2 (*Vglut2)* from Kiss1^ARC^ neurons replicated the obesogenic effect. Furthermore, deletion of the melanocortin 4 receptor (MC4R) from Kiss1 neurons resulted in obesity, reduced energy expenditure and impaired thermogenesis. Optogenetic stimulation of pro-opiomelanocortin (POMC) fibers evoked inward currents in Kiss1 neurons, that were attenuated by MC4R antagonism. Our findings reveal a previously unrecognized neural circuit that mediates melanocortin action on energy expenditure, offering new insights into central mechanisms of metabolic control.

## INTRODUCTION

Kisspeptins, products of the *Kiss1* gene, have emerged as critical regulators at the intersection of reproduction and metabolism. Originally identified for their essential role in reproductive function, kisspeptins are produced primarily by glutamatergic neurons in the arcuate nucleus (ARC) and GABAergic neurons in the anteroventral periventricular nucleus (AVPV/PeN) ^1^. These Kiss1 neurons serve as nodal regulatory centers controlling the patterned release of gonadotropin-releasing hormone (GnRH) from GnRH neurons in the preoptic area of the hypothalamus.

While the reproductive functions of Kiss1 neurons have been extensively characterized ^1^, growing evidence suggests they also play a significant role in metabolic regulation. Several observations support this role: fasting suppresses *Kiss1* expression in rodents and primates ^2–4^; Kiss1 neurons co-express leptin and insulin receptors, indicating they function as metabolic sensors ^5–7^; female Kiss1r null (Kiss1rKO) mice develop marked obesity compared to controls ^8^; and arcuate Kiss1 (Kiss1^ARC^) neurons receive direct inputs from both Agouti-related peptide (AgRP) and pro-opiomelanocortin (POMC) neurons regulating energy homeostasis ^9,10^.

The specific effects of kisspeptin on metabolism, particularly food intake, remain contested. Inhibitory effects on food intake have been reported after central injections of kisspeptin in mice ^11^, jerboa ^12^ and rats ^13,14^, while other studies in humans ^15^, rats ^2,16^, and Kiss1rKO mice ^8^ do not support a significant role of kisspeptin on feeding. Rather, these findings suggest kisspeptin may primarily regulate energy expenditure (EE). This function appears to be mediated specifically by Kiss1^ARC^ neurons, as Kiss1^AVPV/PeN^ neurons are nearly absent in males ^17^ and silencing of Kiss1^ARC^ neurons in female mice increases body weight ^18^. Interestingly, the more severe metabolic phenotype observed after silencing Kiss1^ARC^ neurons compared to the phenotype of Kiss1rKO mice suggests that additional factors from Kiss1 neurons (beyond kisspeptin itself) may contribute to its metabolic regulation.

EE plays a critical role in maintaining overall homeostasis, occurring either through increased physical activity that elevates muscle temperature or through brown adipose tissue (BAT) thermogenesis. In the hypothalamus, α-melanocyte-stimulating hormone (α-MSH), the primary product of POMC neurons, activates the melanocortin 4 receptor (MC4R) to promote satiety and increase EE ^19^, while AgRP antagonizes this receptor to produce opposite effects ^20^. How Kiss1^ARC^ neurons participate in this pathway remains unknown.

While most of α-MSH’s known metabolic actions occur on glutamatergic MC4R-expressing neurons in the paraventricular hypothalamus (PVH), studies have revealed that specific re-introduction of MC4R in PVH neurons restores food intake in MC4RKO mice but fails to recover EE, resulting in only partial (∼60%) body weight normalization ^21^. Similarly, selective deletion of MC4R from PVH neurons produces obesity, though to a lesser degree than when MC4R is ablated from all glutamatergic neurons ^22^. These findings suggest that additional glutamatergic neuronal populations in other brain regions mediate the melanocortin control of energy expenditure. In this context, the potential role of glutamatergic Kiss1^ARC^ neurons in melanocortin signaling presents an intriguing possibility, particularly given that axonal projections from POMC neurons to Kiss1 neurons have been identified ^10^, and we and others have previously shown that Kiss1^ARC^ neurons are direct targets of POMC neurons through MC4R signaling ^23–25^. Whether POMC projections form functional synaptic contacts capable of modulating Kiss1 neuronal activity, potentially positioning Kiss1 neurons as mediators of melanocortin effects on energy expenditure, remains to be explored.

Here, we set out to investigate the role and mechanism of action of Kiss1^ARC^ neurons in energy balance, as well as their integration within the melanocortin system. To this end, we utilized transgenic mouse models to target Kiss1^ARC^ neurons that were ablated or manipulated through chemo- and optogenetics. Furthermore, we removed MC4R from Kiss1 neurons and characterized the metabolic phenotype along with the biophysical properties of these neurons in response to melanocortins and photo-stimulation of POMC neurons. We found that Kiss1^ARC^ neurons regulate energy expenditure through, at least in part, glutamatergic projections to DMH leptin-receptor (Lepr) responsive neurons to regulate the activation of BAT. These neurons are activated by POMC neurons, thus supporting their role as part of the melanocortin machinery that regulates energy expenditure.

## RESULTS

### Kiss1^ARC^ neurons modulate energy status

To evaluate the metabolic function of Kiss1^ARC^ neurons, we bilaterally injected a Cre-dependent AAV-taCasp3 into the ARC of adult (∼3 months old) Kiss1^cre/+^ mice to ablate these neurons (**Figure 1A-C**). This ablation led to a significant increase in body weight (BW) in Kiss1^ARC^ neuron-ablated mice compared to controls, starting from 8 weeks post AAV-taCasp3 injections until at least 12 weeks post-ablation (**Figure 1D**). To assess whether ablation of Kiss1^ARC^ impacts the reproductive axis and induces hypogonadism associated obesity, an additional group of castrated mice injected with a control AAV-mCherry was included. BW in this group did not differ from intact controls (data not shown). Next, to determine whether Kiss1^ARC^ neurons influence food intake, we chemogenetically activated them using a Cre-dependent AAV containing an excitatory chemogenetic receptor construct (AAV-hM3Dq) (**Figure 1E, F**). Animals were fed *ad libitum*, and feeding was monitored 4 hours before (baseline) and 4 hours after clozapine-N-oxide (CNO, 3mg) administration. We observed no changes in food intake following CNO administration (**Figure 1G, H**), in line with previous observations showing that neither chemogenetic activation ^26^ nor viral silencing ^18^ of Kiss1^ARC^ neurons altered food intake. Therefore, we set out to investigate whether the changes in BW could be due to changes in EE. To this end, temperature transponders were implanted subcutaneously above the intrascapular BAT (iBAT) pad to assess thermogenesis after a similar chemogenetic activation of Kiss1^ARC^ neurons (**Figure 1I**), which revealed a significant temperature increase in the iBAT (**Figure 1J**). Of note, BAT thermogenesis lasted for at least 120 min following Kiss1^ARC^ neuron activation with CNO, with a greater magnitude of response than the effect of CNO in controls, suggesting that BAT activation was sustained and it was not caused by the stress from handling. These results demonstrate that Kiss1^ARC^ neurons are capable of modulating energy expenditure through BAT activation.

**Figure 1.**
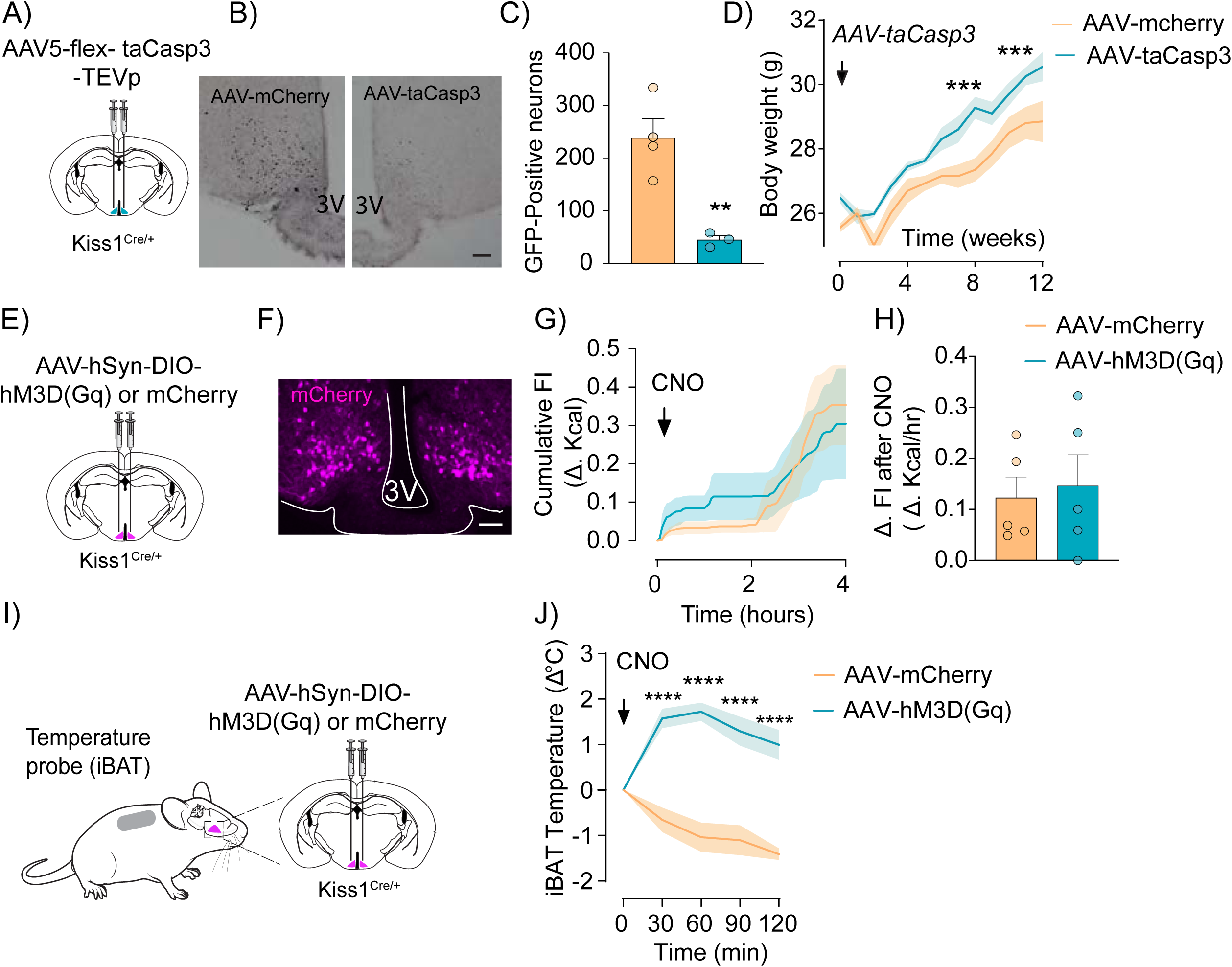
Kiss1^ARC^ neurons modulate energy status. **(A)** Schematic of AAV5-flex-taCasp3-TEVp injection in the ARC of Kiss1^Cre/+^ mice to induce Kiss1 neuron ablation. **(B)** Images of AAV5-taCasp3 injected mice along with their AAV-mCherry injected controls showing GFP chromogenic staining. Scale bar = 100 µm. **(C)** Quantification of GFP positive neurons in the ARC showed a significant decrease in the number of Kiss1 neurons in the ARC in the AAV5-taCasp3 mice (45±7.81, n=3) compared to AAV-mCherry controls (238.5±36.52, n=4). Student’s t test: p= 0.0068, t=4. **p < 0.01. **(D)** Body weight of adult Kiss1^Cre/+^ males from the time of viral injection (Time 0), until 12 weeks post-injections. Two-way ANOVA: n =4/group. 4 weeks post-injection: p=0.1222. 8 weeks post-injection: ***p=0.0001. 12 weeks post-injection: ***p=0.0007. ***p < 0.001. **(E)** Schematic of an AV9-hSyn-DIO-hM3D(Gq)-mCherry injection in the ARC. **(F)** Representative photomicrographs depicting the expression of the AAV9-hM3D(Gq)-mCherry in the ARC. 3V= third ventricle. Scale bar = 100 µm. **(G)** Tracing of cumulative food intake (FI) representing measures every 3 min for 4 hours. Two-way ANOVA. **(H)** The change in cumulative food intake (Δ. FI) from baseline (CNO injection) between AAV-mCherry (0.123±0.040) and AAV-hM3D(Gq) (0.146±0.060), n=5/group. Student’s t test p=0.755, t=0.3230. p>0.05. **(I)** Schematic of iBAT temperature recording of a Kiss1^Cre/+^ mouse injected with an AAV9-hSyn-DIO-hM3D(Gq)-mCherry in the ARC. **(J)** Effect of CNO–hM3Dq stimulation of Kiss1^ARC^ neurons on iBAT temperature. Repeated measures Two-way ANOVA followed by Sidak’s multiple comparisons test: n=4/group. Time 0 is the time of CNO injection: p >0.9999, t=0. 30 min: ****p<0.0001, t= 6.496; 60 min: ****p <0.0001, t= 8.033; 90 min: ****p <0.0001, t= 6.977, 120 min: ****p <0.0001, t= 7.006. Data are represented as mean ± SEM.

### Identification of the Kiss1^ARC^ to BAT activation pathway

To identify the potential pathway by which Kiss1^ARC^ neurons produce an increase in BAT activation, we examined whether the dorsomedial hypothalamus (DMH) neurons were recruited after chemogenetic activation of the Kiss1^ARC^ neurons. To address this, we measured cFos expression in the DMH region, observing a statistically significant increase in cFos levels in the DMH due to CNO-induced activation of the Kiss1^ARC^ neurons (**Figure 2A-D**). Then, we determined whether Kiss1^ARC^ neurons directly innervate the DMH. For this, we evaluated the neuroanatomical projections from Kiss1^ARC^ neurons by using a Cre-dependent AAV opsin AAV-eOPN3. As a result of mapping the potentially target areas of Kiss1^ARC^ neurons (**Figure S1**), we observed Kiss1^ARC^ fibers projecting to the dorsomedial hypothalamus (DMH) (**Figure 2C, Figure S1I, J**), which is known to gate the activation of BAT ^27,28^. This suggests that Kiss1^ARC^ neurons directly contact and activate DMH neurons. Next, we investigated whether the DMH neurons receiving inputs from Kiss1^ARC^ neurons project to the raphe pallidus (RPa), a key relay linking the DMH to the BAT ^27,28^. To assess this pathway, we injected retrobeads into the RPa and examined the DMH for retrogradely beads-labeled neurons contacted by Kiss1^ARC^ projections. We found that Kiss1^ARC^ fibers were in close apposition to RPa-projecting (retrobeads-labeled) neurons in the DMH (**Figure 2E, F**). The DMH neurons projecting to the RPa have been previously identified as Lepr/CART-expressing neurons. To reveal the identity of DMH neurons receiving input from Kiss1^ARC^ neurons, we co-stained for cFos and STAT3 following chemogenetic activation of Kiss1^ARC^ neurons with CNO, combined with leptin administration to label leptin-responsive neurons. Clear co-localization of cFos and STAT3 was detected in the DMH (**Figure 2G-I**), indicating activation of leptin-responsive neurons by Kiss1^ARC^ stimulation and further supporting the role of Kiss1^ARC^ neurons as upstream regulators of this thermogenic pathway.

**Figure 2.**
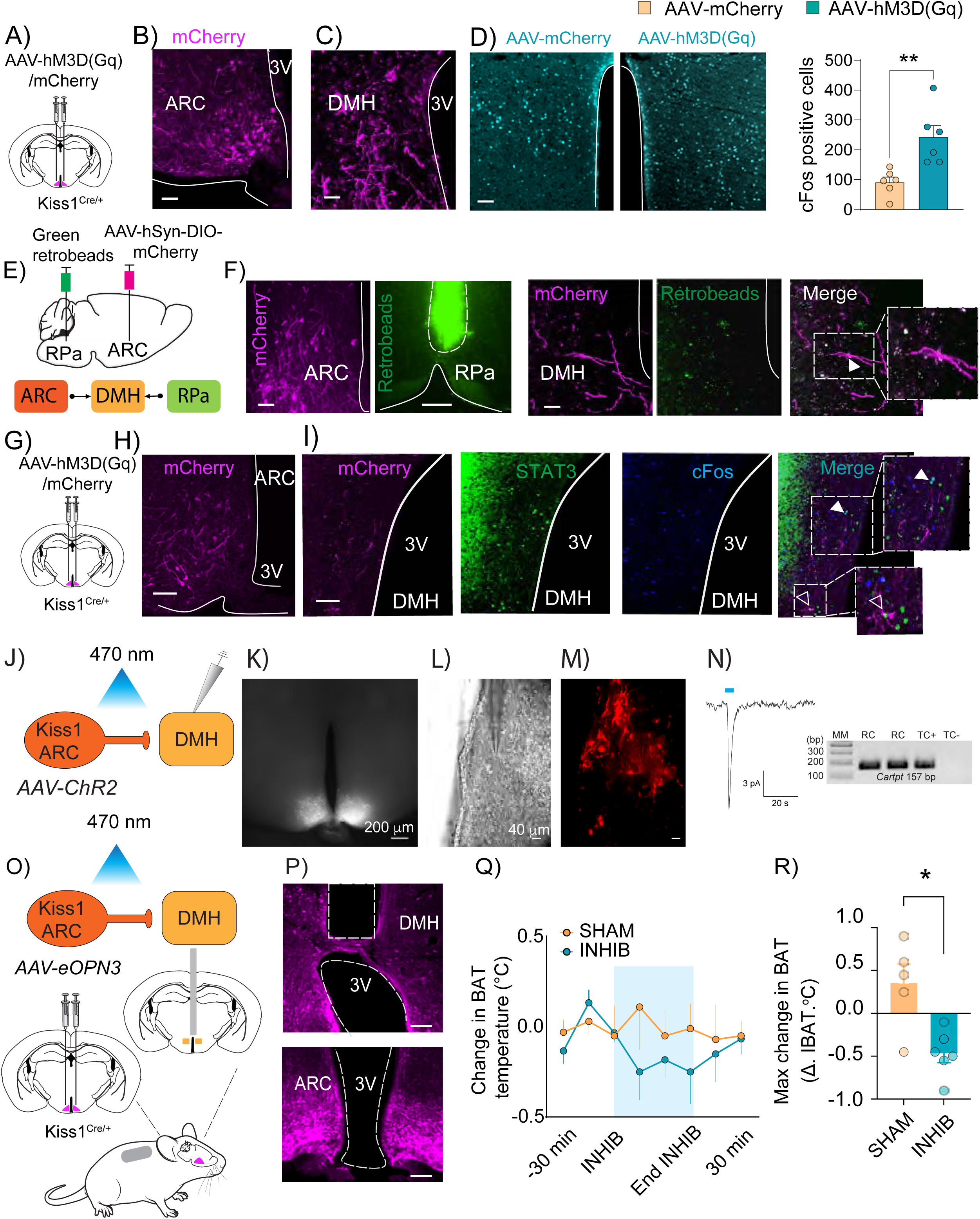
Kiss1^ARC^ neurons activate Cart/Lepr neurons in the DMH to regulate BAT thermogenesis. **(A)** Schematic of AAV9-hSyn-DIO-hM3D(Gq)-mCherry injection in the ARC of Kiss1^Cre/+^ mice. **(B)** photomicrographs depicting the expression of the AAV9-hM3D(Gq)-mCherry in the ARC. 3V= third ventricle. Scale bar = 100µm. **(C)** Representative expression of Kiss1^ARC^ neurons projections in the DMH. **(D)** Representative expression of cFos in AAV-hM3D(Gq) vs AAV-mCherry mice (Scale bar = 100 µm) and quantification of cFos positive neurons in the DMH showing a significant increase in cFos expression in the AAV-hM3D(Gq) group (242.2±38.80) vs the AAV-mCherry (91.17±17.58) group. Student’s t test: **p=0.0053, t=3. n=6/group. **(E)** Schematic of AAV9-hSyn-DIO-hM3D(Gq)-mCherry injection in the ARC and green retrobeads in the RPa of Kiss1^Cre/+^ mice (n=3) to map the ARC-DMH-RPa pathway. **(F)** Representative photomicrographs depicting mCherry expression in Kiss1^ARC^ neurons, retrobeads expression in the RPa, and Kiss1 axonal projections contacting retrobeads positive neurons in the DMH. Inset shows a zoom in of the boxed region, illustrating contact and close apposition of Kiss1 fibers and RPa projecting neurons. Scale bar = 100 µm. **G)** Schematic of AAV9-hSyn-DIO-hM3D(Gq)-mCherry injection in the ARC in Kiss1^Cre/+^ mice (n=3). **(H)** Representative photomicrographs depicting mCherry expression in Kiss1^ARC^ neurons (Scale bar 100 µm), projecting to **(I)** the DMH, where they form close apposition with STAT3 (Lepr expressing) neurons. The insets provide a zoom in of the boxed regions highlighting two populations of STAT3 neurons: those contacted by Kiss1 fibers (empty arrows) and those that are cFos positive (full arrows). Scale bar = 100 µm. **(J)** Schematic of the *in vitro* optogenetic stimulation of Kiss1^ARC^ fibers in the DMH. Viral injection of AAV-ChR2 was used to drive expression of excitatory opsin (ChR2) and mCherry in Kiss1^ARC^ neurons. Recordings were made in DMH neurons in response to optogenetic stimulation. (**K)** Widefield image of Kiss1^ARC^ mCherry expression. Scale bar = 200 µm. **(L)** High-power brightfield image of the DMH near the 3V. Scale bar = 40 μm. **(M)** Fluorescent widefield image of the same region from **(K)**. **(N)** Postsynaptic response in DMH neuron following optogenetic stimulation of Kiss1^ARC^ neurons. Cytosol of recorded neurons was harvested, underwent RT-PCR, and tested for expression of Cart. (See also Figure S5). **(O)** Schematic of the *in vivo* optogenetic stimulation of Kiss1^ARC^ fibers in the DMH in Kiss1^Cre/+^ mice implanted with temperature transponders above the iBAT pad. Viral injection of AAV-eOPN3 was used to drive expression of inhibitory opsin (eOPN3) and mCherry in Kiss1^ARC^ neurons. **(P)** Representative expression of AAV-eOPN3 in the ARC and location of optic fiber above the DMH. Scale bar 100 µm. **(Q)** The iBAT temperature was recorded 30 min before, 1 hour during and 30 min after photoinhibition (INHIB). **(R)** The mean temperature change (Δ. iBAT) during the STIM period (1 hour) was significantly different between SHAM (-0.01500±0.02096, n=5) and INHIB (-0.1167±0.04499, n=6) mice. Student’s t test: *p=0.0454, t=1.960. Blue transparent area represents STIM period. See figure S1 for the mapping of Kiss1 neurons projections in the male mouse brain.

We have previously shown in females that Kiss1^ARC^ neurons make direct glutamatergic projections to DMH neurons expressing *Cart* and *Lepr* ^29^. To confirm that this is also true in males, bilateral injections of AAV1-DIO-mCh-ChR2 were made into the ARC of Kiss1^cre/+^ mice (**Figure 2J, K**). Cells were patched in the DMH where mCherry fibers were present (**Figure 2L, M**). Voltage clamp recordings were made during brief (5 ms) blue light pulses. Postsynaptic currents were identified based on the clear delay between stimulation and response (**Figure 2N**). Once an excitatory postsynaptic current was observed, the location of the electrode was noted and the cytosol of the cell was harvested. Harvested cells underwent RT-PCR and were tested for the presence of *Cart* as marker of Lepr/CART+ neurons. We were able to detect *Cart* mRNA in three out of nine cells (**Figure 2N**). These findings further confirm the Kiss1^ARC^ to DMH circuits and demonstrate the existence of postsynaptic glutamatergic connections, as previously shown in females ^29^. However, the presence of synaptic connections with Lepr-expressing DMH neurons does not definitively prove that Kiss1^ARC^ neurons regulate BAT exclusively through this pathway, given that other hypothalamic areas receiving projections from Kiss1^ARC^ neurons could also contribute to this action (**Figure S1**). Thus, to directly test the functional relevance of the Kiss1^ARC^ to DMH projections, we optogenetically inhibited Kiss1^ARC^ neuron terminals in the DMH by injecting a Cre-dependent inhibitory opsin in the ARC (AAV-eOPN3, an inhibitory opsin that is suitable for long-lasting inhibition of cell bodies or synaptic terminals using low-power (blue laser, 470 nm) illumination ^30^. This approach allowed us to inhibit the Kiss1^ARC^ DMH pathway while simultaneously recording BAT temperature by using a thermography method ^31,32^ (**Figure 2O, P**). Moreover, our optogenetic inhibition method prevented the potential confounding effect from the collateral activation of other brain areas after Kiss1^ARC^ neuron activation. During a one-hour photoinhibition period, AAV-eOPN3 induced suppression of BAT thermogenesis and a consequent reduction of BAT temperature (**Figure 2Q, R**). These findings demonstrate that Kiss1^ARC^ neurons promote EE through the activation of the CART/Lepr^DMH^ neuron to RPa circuit.

### Kiss1^ARC^ neurons activate the DMH–RPa–BAT pathway via glutamate

The glutamatergic activation of Lepr/CART^DMH^ neurons by photo-stimulation of Kiss1^ARC^ terminals **(Figure 2N)** suggested a potential kisspeptin-independent mechanism. To directly test this, we chemogenetically activated Kiss1^ARC^ neurons of Kiss1^Cre/Cre^ mice -which lack functional kisspeptin (Kiss1KO)-as in **Figure 1**. Kiss1^ARC^ neuron terminals were observed in the DMH of Kiss1KO mice and CNO administration in AAV-hM3D(Gq) injected mice significantly increased cFos expression in the DMH (**Figure 3A-D**). Importantly, this activation also correlated with a rise in iBAT temperature in Kiss1KO mice starting at 30 min and persisting for at least 120 min following Kiss1^ARC^ activation (**Figure 3E-G**). Next, we hypothesized that if the change in BW observed after Kiss1^ARC^ neuron ablation (**Figure 1**) is driven by the loss in glutamatergic signaling, then selective ablation of glutamate release from these neurons should reproduce the increase in BW. To test this, we used a Cre-dependent CRISPR virus (AAV1-FLEX Sacas9-SgSlc17a6) ^33,34^ to delete *Vglut2* selectively from Kiss1^ARC^ neurons of adult Kiss1^cre/+^ mice fed a 60% high fat diet (HFD) (**Figure 3H, I**). Kiss1-Vglut2KD mice displayed a significant increase in BW starting 3 weeks after the onset of HFD exposure, which persisted through at least 8 weeks (**Figure 3J**). This phenotype closely resembled that of mice with ta-Casp3-mediated ablation of Kiss1^ARC^ neurons (**Figure 1D),** supporting a key role for glutamatergic signaling from Kiss1^ARC^ neurons in regulating EE.

**Figure 3.**
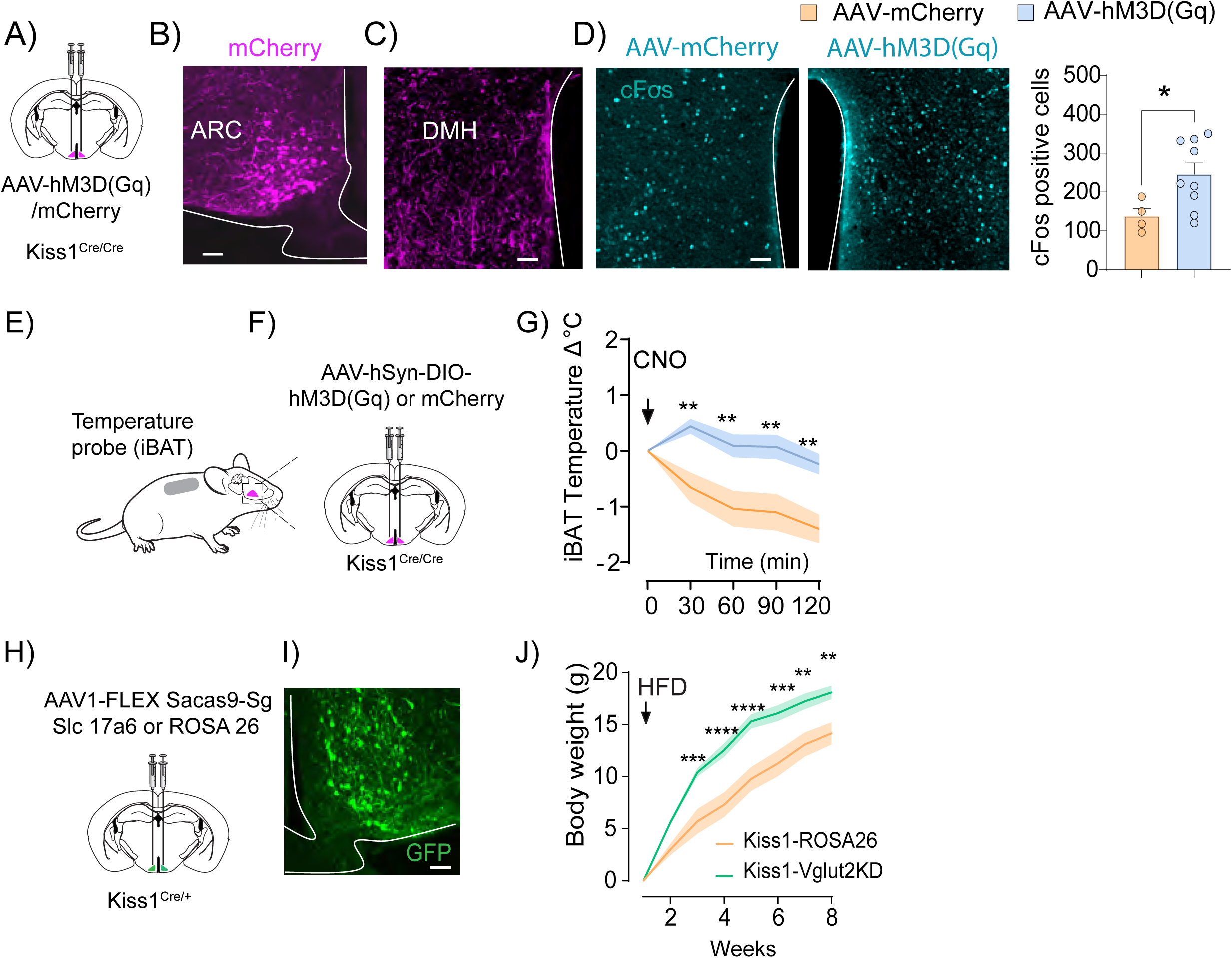
Activation of Kiss1^ARC^ neurons in Kiss1KO mice activates DMH neurons and BAT thermogenesis, and Vglut2 deletion in Kiss1^ARC^ induces obesity. **(A)** Schematic of AAV9-hSyn-DIO-hM3D(Gq)-mCherry injection in the ARC of Kiss1^Cre/Cre^ (Kiss1KO) mice. **(B)** Representative photomicrographs depicting the expression of the AAV9-hM3D(Gq)-mCherry in the ARC. Scale bar = 100 µm. **(C)** Representative expression of Kiss1^ARC^ neurons projections in the DMH of Kiss1KO mice. **(D)** Representative expression of cFos in AAV-hM3D(Gq) vs AAV-mCherry mice) and quantification of cFos positive neurons in the DMH showing a significant increase in cFos expression in the AAV-hM3D(Gq) group (245.7 ± 29.22, n=9) vs the AAV-mCherry (138.0 ± 20.02, n=4) group. Student’s t test: *p=0.0414, t=2.30. Scale bar = 100 µm. **(E)** Schematic of iBAT temperature recording of a Kiss1KO mouse injected with an AAV9-hSyn-DIO-hM3D(Gq)-mCherry in the ARC **(F)**. **(G)** Effect of CNO–hM3Dq stimulation of Kiss1^ARC^ neurons on iBAT temperature. Repeated measures Two-way ANOVA followed by Sidak’s multiple comparisons test. Time 0 is the time of CNO injection: p >0.9999, t=0. 30 min: **p=0.0036, t=3.579; 60 min: **p=0.0026, t=3.681; 90 min: **p=0.0016, t=3.832, 120 min: **p =0.0018, t=3.796. **(H)** Schematic of AAV1-FLEX Sacas9-SgSlc17a6 injection in the ARC of Kiss1^Cre/+^ mice. **(I)** Representative photomicrographs depicting the expression of the CRISPR AAV1-FLEX KASH EGFP in the ARC to specifically delete *Vglut2* from Kiss1^ARC^ neurons. Scale bar = 100 µm. **(J)** Body weight gain of adult Kiss1-Vglut2KD and control Kiss1-ROSA26 males following submission to a 60% HFD. Repeated measures Two-way ANOVA followed by Sidak’s multiple comparisons test. Week 3: ***p=0.0005, t=4.323. Week 4: ****p<0.0001, t=4.822. Week 5: ****p<0.0001, t=5.082. Week 6: ***p=0.0003, t=4.438. Week 7: **p=0.0033, t=3.810. Week 8: **p=0.0065, t=3.620.

### Congenital deletion of MC4R from Kiss1 neurons induces obesity

Given the action of Kiss1^ARC^ neurons in the regulation of EE described above, their expression of *Mc4r* ^23–25^ and their reported interactions with the neurons within the melanocortin system, i.e. AgRP and POMC neurons ^24,35,36^, we set out to determine the metabolic consequences of selectively ablating *Mc4r* from Kiss1 neurons in mice. We first confirmed the selective ablation of *Mc4r* from Kiss1 neurons of the arcuate nucleus in Kiss1^MC4RKO^ mice (**Figure S2**), while its expression remained intact in the PVH, the primary site of MC4R action in the control of energy balance (**Figure S2**). Unexpectedly, Kiss1^MC4RKO^ mice exhibited a remarkable increase in BW on regular chow diet, without changes in body length (**Figure 4A-C**). This increase was due to a significant gain in fat mass with a slight reduction in lean mass (**Figure 4D-F**). Supporting this metabolic phenotype, Kiss1^MC4RKO^ mice showed greater hepatic lipid accumulation, indicative of hepatic steatosis (**Figure 4G, H**). These mice also displayed a significant reduction in the uncoupling protein 1 (*Ucp1)* expression in the BAT, consistent with impaired EE (**Figure 4I**). Interestingly, glucose metabolism was preserved in Kiss1^MC4RKO^ mice (**Figure 4J, K)**. To confirm that the removal of MC4R from Kiss1 neurons did not induce hypogonadism, which could affect body weight and composition, we assessed the reproductive phenotype of Kiss1^MC4RKO^ mice and observed no differences in the age of puberty onset, BW at the age of puberty onset (suggesting that the detected BW increase in KOs develops during adulthood as shown in **Figure 4A**), and present normal levels of luteinizing hormone (LH) and testosterone in adulthood (**Figure S3**). Because *Kiss1* is expressed in other tissues beyond the brain, we assessed the expression of *Mc4r* mRNA in the preoptic area (POA), mediobasal hypothalamus (MBH), testis, pituitary, BAT, white adipose tissue (WAT), liver and muscle. *Mc4r* was mostly detected at the central nervous system level, in accordance with its described neuronal effect, and no changes were detected in Kiss1^MC4RKO^ mice compared to controls in any tissue (**Figure S4**), not even in the MBH due to the small relative size of the Kiss1 population.

**Figure 4.**
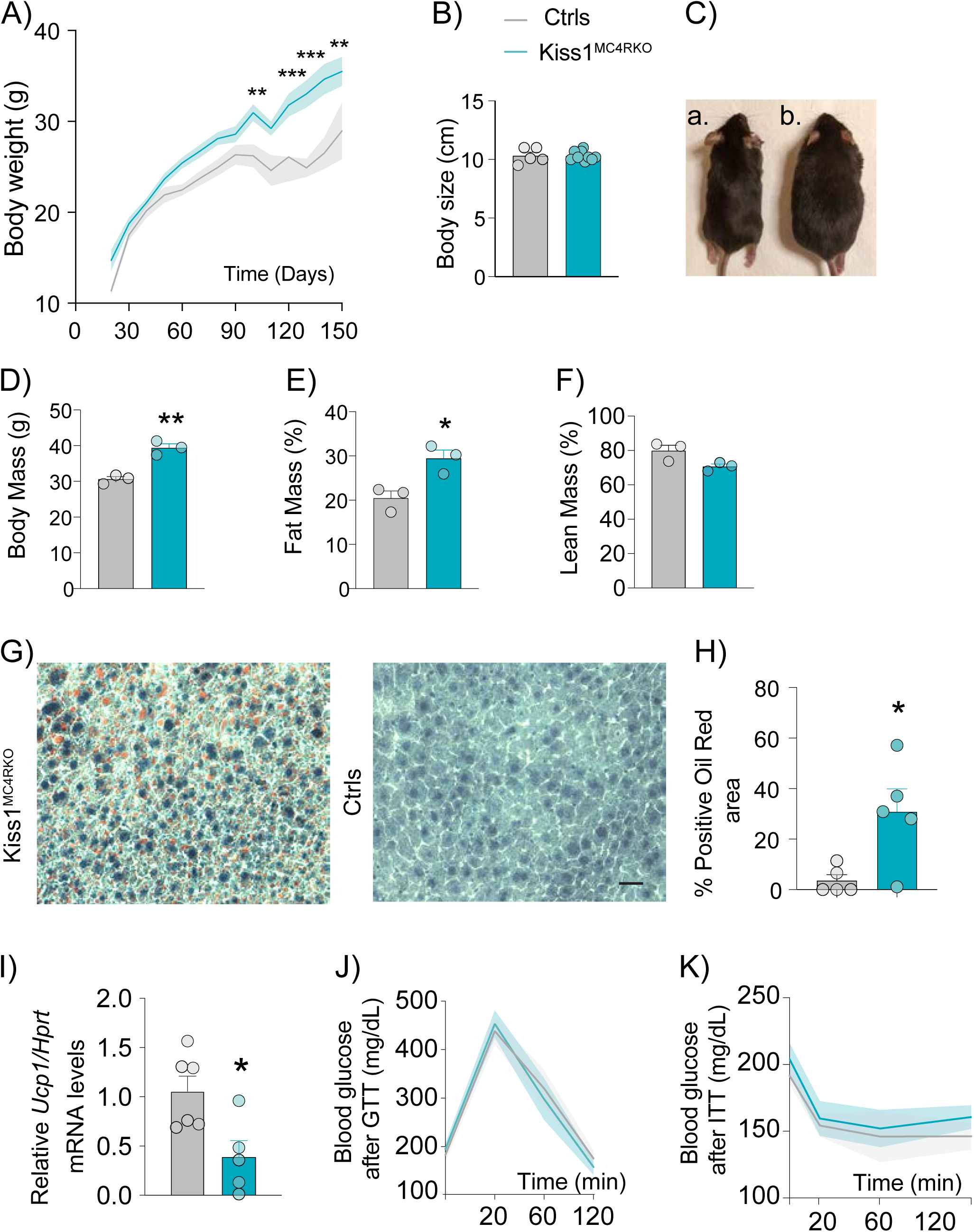
Congenital deletion of MC4R from Kiss1 neurons induces obesity. **(A)** Body weight curve over time showing that Kiss1^MC4RKO^ mice (n=14) gain significantly more weight than their WT littermates (n=6) starting from post-natal day (PND) 100. Significant differences in body weights were found at PND 100: **p=0.0066, t=3.552. PND 130: ***p=0.0002, t=4.514. PND 140: ***p=0.0002, t=4.503. PND 150: **p=0.0058, t=3.590. Data was analyzed using a mixed-effects model for repeated measures to account for missing time points in some animals) followed by Sidak’s multiple comparisons test. **(B)** Body size of adults (6 months) Kiss1^MC4RKO^ mice (10.31 ± 0.1328, n=9) and their WT littermates (10.32 ± 0.295, n=5) showing similar body length between the two groups. Student’s t test: p=0.975. **(C)** Representative images showing similar body length and different body weight between an adult Kiss1^MC4RKO^ mouse (b) and a control littermate mouse (a). Kiss1^MC4RKO^ mice and WT littermates (n=3/group) MRI analysis revealed significant differences in **(D)** Body mass (**p=0.0027. Kiss1^MC4RKO^: 39.40 ± 1.127; WT: 30.63 ± 0.705), **(E)** percentage (%) of fat mass (*p=0.021. Kiss1^MC4RKO^:29.47 ± 1.865; WT: 20.46 ± 1.599), and a tendency to a decrease **(F)** % of lean mass (p=0.548. Kiss1^MC4RKO^: 70.78 ± 1.415; WT: 79.98 ± 3.117). Student’s t test. **(G)** Representative images showing stained lipid droplets using Oil red staining in Kiss1^MC4RKO^ mice and WT littermates (n=5/group). Scale bar is 100 µm. **(H)** Quantification of the stained lipid droplets showing a significant increase in lipid deposition in the liver in Kiss1^MC4RKO^ mice (30.84 ± 9.013) compared to WT controls (3.607 ± 2.331). *p=0.0191. Student’s t test. **(I)** *Ucp1* mRNA expression was significantly decreased in the Kiss1^MC4RKO^ mice (0.389 ± 0.165, n=5) compared to their control littermates (1.056 ± 0.153, n=6). Student’s t test. *p=0.0163. A glucose tolerance test **(K)** revealed normal blood glucose levels between Kiss1^MC4RKO^ mice and their WT littermates (n=8/group). Repeated measures Two-way ANOVA followed by Sidak’s multiple comparisons test. Time 0: p= 0.939. Time 20 min: p= 0.992. Time 60 min: p= 0.992. Time 120 min: p= 0.914. Similarly, an insulin tolerance test **(L)** showed no differences between groups (n=8/group). Repeated measures Two-way ANOVA followed by Sidak’s multiple comparisons test. Time 0: p=0.923. Time 20 min: p= 0.996. Time 60 min: p= 0.994. Time 120 min: p= 0.874. Data are presented as mean ± SEM. *See also Figure S2 for validation of the mouse model, Figure S3 for the reproductive phenotype of these mice, and Figure S4 for the peripheral validation of these mice*.

### Reduced energy expenditure and altered circadian feeding behavior in Kiss1^MC4RKO^ mice

To further characterize their metabolic phenotype, Kiss1^MC4RKO^ mice were placed on a HFD and assessed through a Comprehensive Lab Animal Monitoring System (CLAMS). HFD exacerbated the increase in BW in Kiss1^MC4RKO^ mice (**Figure 5A**). As expected, and consistent with the ablation/activation of Kiss1^ARC^ neurons in **Figure 1**, the cumulative food intake was not altered in Kiss1^MC4RKO^ mice (**Figure 5B**). However, Kiss1^MC4RKO^ mice lost the circadian rhythm of feeding behavior, leading to the same food consumption during the light and dark phases, in stark contrast to WT mice that predominantly ate during the dark phase (**Figure 5C, D**). This disruption in rhythmic feeding correlated with altered feeding bouts, indicating that the amount of food eaten per bout was similar between the dark and light phases in the KO mice (**Figure 5E, F**). This change in circadian feeding behavior is in line with previous studies in Kiss1^ARC^ neuron-silenced female mice ^18^. Interestingly, water consumption mirrored the alterations in feeding behavior, with Kiss1^MC4RKO^ mice exhibiting a loss of the normal circadian rhythm of drinking behavior (**Figure 5G, H**). Analysis of VO2 consumption and VCO2 production evidenced a decrease in EE that is mostly evident during the dark phase (**Figure 5I-L**). This decrease was not associated to changes in locomotor activity (**Figure 5M, N**). Consistent with these findings, heat production also failed to increase during the dark phase in Kiss1^MC4RKO^ mice, in contrast to controls (**Figure 5O, P**), further supporting impaired regulation of energy expenditure. While control mice did not exhibit the expected increase in RER during the dark phase (**Figure 5Q, S**), Kiss1^MC4RKO^ mice showed a significant decrease in RER during the dark phase. This is likely attributable to the impaired feeding behavior during the dark phase, leading mice to rely more heavily on fat oxidation rather than carbohydrate utilization. Together, these findings support a critical role of MC4R signaling in Kiss1 neurons in the regulation of EE.

**Figure 5.**
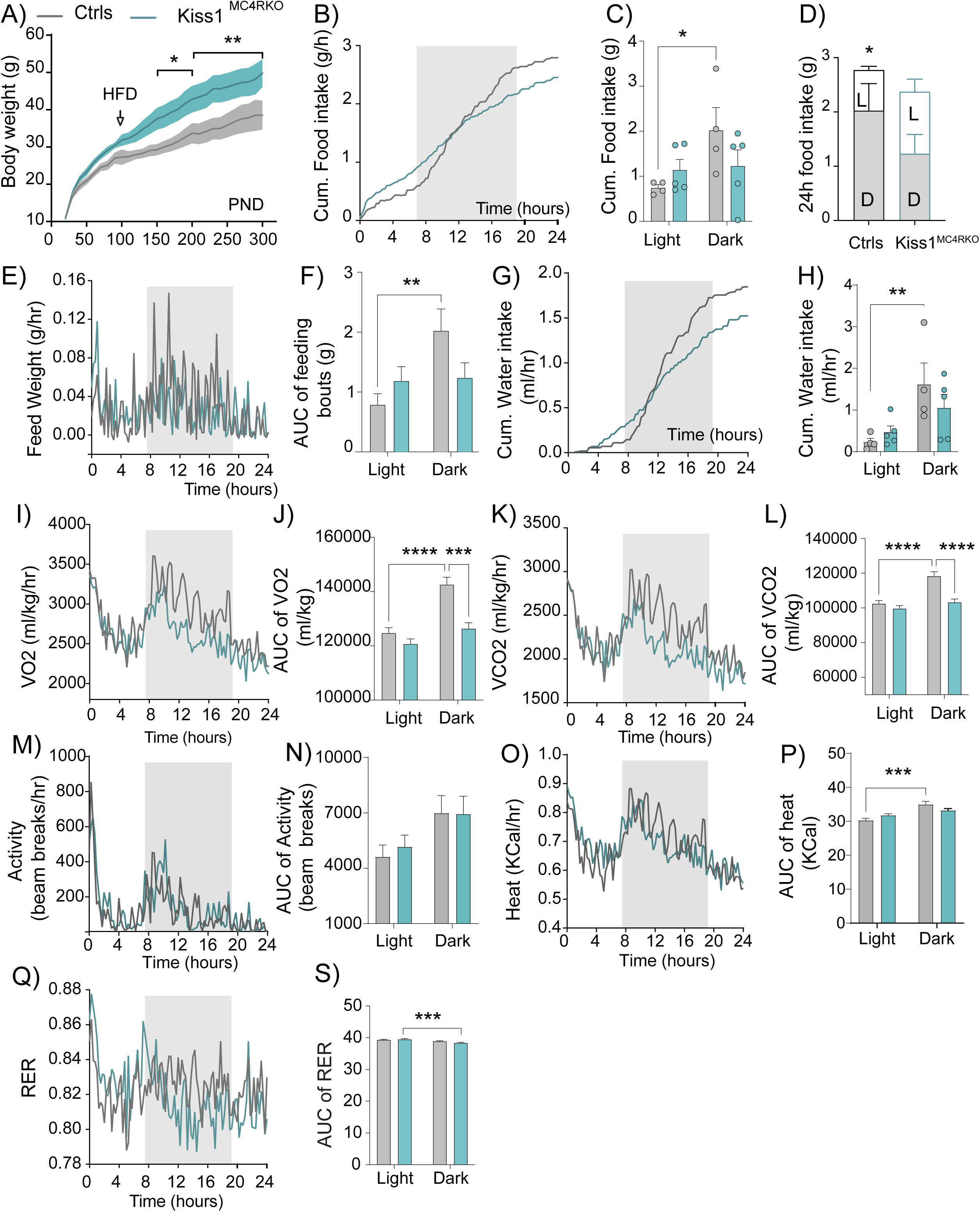
High fat diet exacerbates obesity in Kiss1^MC4RKO^ mice and reveals impaired energy expenditure and circadian rhythmicity of feeding and drinking behaviors. **(A)** Body weight curve over time of Kiss1^MC4RKO^ mice and their WT littermates. BW was monitored from weaning, and at post-natal day (PND 90) animals were submitted to 60% high fat diet (HFD) and body weight was monitored until PND 300. HFD exacerbated the obese phenotype in Kiss1^MC4RKO^ mice. Significant differences in body weights were found at PND 150: *p=0.0232, t=2.288. PND 160: *p=0.0252, t=2.246. PND 170: ***p=0.0182, t=.2.380 PND 180: **p=0.0189, t=2.366. PND190: *p=0.0136, t=2.490. PND 200: *p=0.0131, t=2.504. PND 210: **p=0.0073, t=2.712. PND 220: **p=0.0071, t=2.721. PND 230: **p=0.0038, t=2.932. PND 240: **p=0.0039, t=2.922. PND 250: **p=0.0053, t=2.817. PND 260: **p=0.0067, t=2.740. PND 270: **p=0.0079, t=2.684. PND 280: **p=0.0088, t=2.644. PND 290: **p=0.0084, t=2.662. PND 300: **p=0.0023, t=3.086. Data was analyzed using a mixed-effects model for repeated measures to account for missing time points in some animals) followed by Sidak’s multiple comparisons test. **(B)** Waveform of cumulative food intake curve over 24 hours showing feeding behavior of Kiss1^MC4RKO^ mice (n=5) and their WT littermates (n=4) during light and dark (grey area) phases. **(C)** Mean of cumulative food intake per mouse during light and dark phases in WT controls (Light: Control vs. Dark: Control: *p=0.0198, t=2.630), and Kiss1^MC4RKO^ mice (Light: Kiss1^MC4RKO^ vs. Dark: Kiss1^MC4RKO^: p=0.838, t=0.2072). Two-way ANOVA. **(D)** Average of 24 hours food intake with Light:Dark-phase distribution. Mixed model analysis: Control Light vs Dark: *p=0.0267, t=2.828; Kiss1^MC4RKO^ Light vs Dark: p=0.974, t=0.207). **(E)** Waveform of feeding bouts curve over 24 hours of Kiss1^MC4RKO^ mice (n=5) and their WT littermates (n=4) during light and dark phases. **(F)** Area under the curve (AUC) of feeding bouts over 24 hours showing significant light-dark phase differences in the control group (Control Light vs Dark: **p=0.0071, t=3.152) as compared to Kiss1^MC4RKO^ mice: p=0.8821, t=0.1511). Two-way ANOVA. **(G)** Waveform of cumulative water intake curve over 24 hours showing drinking behavior of Kiss1^MC4RKO^ mice (n=5) and their WT littermates (n=4) during light and dark phases. **(H)** Mean of cumulative water intake showing expected differences in drinking behavior between light and dark phases in the WT controls (Light:Control vs. Dark:Control: **p=0.0086, t=3.054), while these differences are impaired in the Kiss1^MC4RKO^ mice (Light: Kiss1^MC4RKO^ vs. Dark: Kiss1^MC4RKO^: p=0.172, t=1.436). Two-way ANOVA. **(I)** Waveform of oxygen consumption (VO2) during 24 hours period of Kiss1^MC4RKO^ mice (n=5) and their WT littermates (n=4) during light and dark phases. **(J)** AUC of VO2 consumption over 24 hours showing impaired circadian rhythmicity of VO2 in Kiss1^MC4RKO^ (Light: Kiss1^MC4RKO^ vs. Dark: Kiss1^MC4RKO^: p=0.067, t= 1.983) compared to controls (Light:Control vs. Dark:Control: ****p<0.0001, t= 5.597), in addition to a significant decrease of VO2 in Kiss1^MC4RKO^ mice compared to controls during the dark phase (Dark Ctrls vs Kiss1^MC4RKO^: ***p=0.0001, t=5.349). Two-way ANOVA. **(K)** Waveform of Carbone dioxide production (CO2) during 24 hours period of Kiss1^MC4RKO^ mice (n=5) and their WT littermates (n=4) during light and dark phases. **(L)** AUC of VCO2 production over 24 hours showing impaired circadian rhythmicity of VCO2 in Kiss1^MC4RKO^ (Light: Kiss1^MC4RKO^ vs. Dark: Kiss1^MC4RKO^: p=0.180, t= 1.409) compared to controls (Light: Control vs. Dark: Control: ****p<0.0001, t= 5.502), in addition to a significant decrease of VO2 in Kiss1^MC4RKO^ mice compared to controls during the dark phase (Dark Ctrls vs Kiss1^MC4RKO^: ****p<0.0001, t=5.508). Two-way ANOVA. **(M)** Waveform of total physical activity as measured via beam breaks activity of Kiss1^MC4RKO^ mice (n=5) and their WT littermates (n=4) during light and dark phases. **(N)** AUC of physical activity over 24 hours showing no significant differences between Kiss1^MC4RKO^ mice (Light: Kiss1^MC4RKO^ vs. Dark: Kiss1^MC4RKO^: p= 0.128, t=1.617) and their controls (Light: Control vs. Dark: Control: p=0.075, t=1.924). No significant differences were found between the two groups in light (Light Ctrls vs Kiss1^MC4RKO^: p=0.649, t=0.463) or dark (Dark Ctrls vs Kiss1^MC4RKO^: p=0.969, t=0.039) phases. Two-way ANOVA. **(O)** Waveform of total heat production in Kiss1^MC4RKO^ mice (n=5) and their WT littermates (n=4) during light and dark phases. **(P)** AUC of heat production over 24 hours showing the expected increase in heat in WT mice during the dark phase (Light: Control vs. Dark: Control: ***p=0.0002, t=4.999) that is absent in the Kiss1^MC4RKO^ mice (Light: Kiss1^MC4RKO^ vs. Dark: Kiss1^MC4RKO^: p=0.103, t=1.739). There is a tendency for a decrease in heat production in Kiss1^MC4RKO^ mice compared to controls during the dark phase, but it doesn’t reach statistical significance (Dark Ctrls vs Kiss1^MC4RKO^: p=0.070, t=1.956). Two-way ANOVA. **(Q)** Waveform of Respiratory Exchange Ratio (RER) in Kiss1^MC4RKO^ mice (n=5) and their WT littermates (n=4) during light and dark phases. **(S)** AUC of RER across light and dark phases in Kiss1^MC4RKO^ mice and controls over 24 hours. Control mice did not show a significant change in RER between light and dark phases (Light:Control vs. Dark:Control: p=0.115, t= 1.680), whereas Kiss1^MC4RKO^ mice exhibit a significant decrease during the dark phase (Light:Kiss1^MC4RKO^ vs. Dark:Kiss1^MC4RKO^: ***p=0.0003, t=4.737) indicating altered substrate utilization. Two-way ANOVA. Data are presented as mean ± SEM.

### POMC neurons synapse on and excite Kiss1^ARC^ neurons via glutamate and αMSH release

Kiss1^ARC^ neurons are known to send glutamatergic projections to both NPY/AgRP and POMC neurons ^37^. Though POMC neurons have been shown to make reciprocal contact with NPY/AgRP neurons ^38–40^, and whether POMC neurons also synapse on Kiss1^ARC^ neurons in the male mouse has not been described. Yet, the expression of MC4Rs in Kiss1^ARC^ neurons ^23–25^ would suggest that such connections exist. To test this hypothesis, we used brain slices from intact POMC-Cre male mice (>2 months) that had been injected with an AAV to drive the expression of mCherry and the excitatory opsin channelrhodopsin 2 (ChR2) (**Figure 6A**). For whole-cell patch recordings we targeted non-fluorescent neurons in the ventral ARC that were densely surrounded by POMC fibers (**Figure 6B**), where Kiss1 cell bodies are concentrated (**Figure 6C**). Neurons exhibiting an input resistance of less than 1.0 GΩ ^41,42^, thereby excluding NPY/AgRP neurons, were probed for postsynaptic responses using brief (5 ms) single pulses of 470 nm blue light. With successful bilateral injection/infection, ∼25% of the patched neurons exhibited postsynaptic currents (PSCs) following optogenetic stimulation. The majority (16/20) displayed only a rapid, inward current that was sensitive to the AMPA/Kainate receptor antagonist CNQX (10 μM), indicating ionotropic glutamatergic signaling (**Figure 6D**). The slight delay (2-5 ms) from the optogenetic stimulation and onset of the fast EPSC strongly suggested that there was a postsynaptic response to POMC neurotransmitter release with no intervening cells. However, in order to demonstrate that POMC neurons make direct projections to Kiss1^ARC^ neurons, we used a “rescue” protocol from previous channelrhodopsin assisted circuit mapping studies ^37,40,43,44^. After eliminating the optogenetic response with the addition of tetrodotoxin (TTX, 1 μM) to the bath, we rescued the postsynaptic glutamate response with the addition of the K^+^ channel blockers 4-aminopyridine (4-AP; 0.5 mM) and tetraethylammonium (TEA; 7.5 mM) (**Figure 6E**). This supports a direct synaptic contact between POMC and Kiss1^ARC^ neurons. A small subset of recordings (4/20) suggested a mix of GABA and glutamate release (both inward and outward currents, **Figure 6F**). Kiss1 cells were positively identified by either the presence of a persistent sodium current ^45^ or *posthoc* identification via sc-PCR (**Figure 6G**). Therefore, POMC neurons appear to primarily utilize glutamate for fast, amino acid neurotransmission to Kiss1^ARC^ neurons.

**Figure 6.**
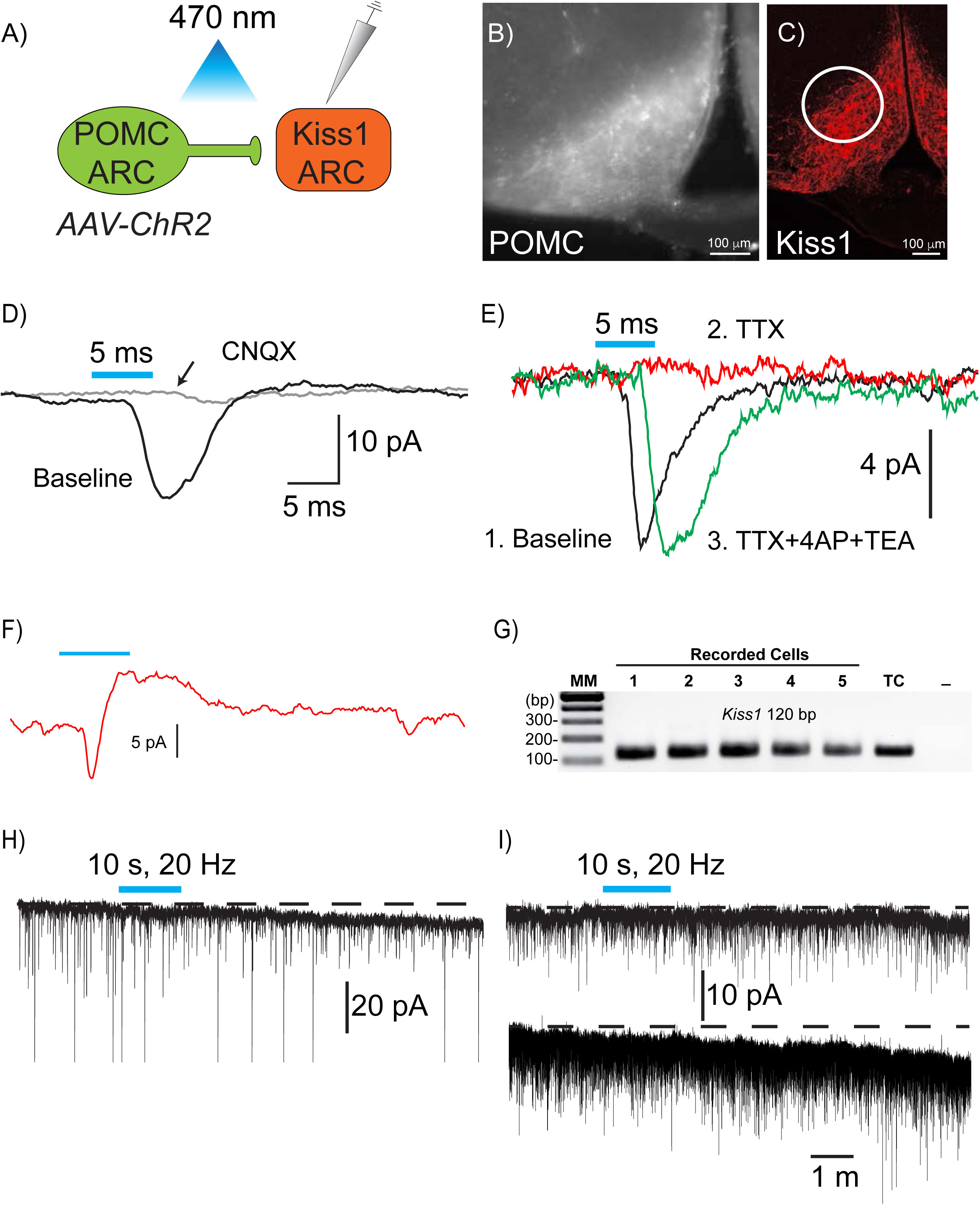
POMC neurons synapse on and excite Kiss1^ARC^ neurons via glutamate and αMSH release. **A)** POMC neurons expressing mCherry-Channelrhodopsin fusion protein were stimulated with blue light while recording from Kiss1^ARC^ neurons. **(B)** Widefield immunofluorescence image of mCherry labeling in a recorded slice. Scale bar = 100 μm. **(C)** Confocal image of mCherry labeling of Kiss1^ARC^ neurons. Scale bar is 100 μm. **(D)** Brief flashes of blue light elicited rapid inward currents blocked by CNQX, indicating AMPA/Kainate-receptor mediated glutamatergic signaling. **(E)** Following TTX, the optogenetically induced response was eliminated. Block of K+ channels lead to a rescue of the response, indicating that POMC neurons make direct synaptic contact with Kiss1 neurons. **(F)** Example of an inward and outward postsynaptic current, suggesting mixed glutamatergic and GABAergic input. **(G)** *Posthoc* identification of recorded cells as Kiss1 neurons. **(H)** Example of a quickly developing inward current following high-frequency optogenetic stimulation. **I)** In some instances, the inward current took several minutes to develop.

Since Kiss1^ARC^ neurons excite POMC neurons through glutamatergic receptors, both ionotropic and metabotropic, and knockout of *Vglut2* eliminates the ability of sex steroids to prevent a conditioned place preference to sucrose ^37^, these two ARC subpopulations may collaborate to maintain energy homeostasis. Furthermore, based on the presence of MC4R expression in Kiss1^ARC^ neurons ^23–25^, we hypothesized that high frequency optogenetic stimulation would elicit peptide release ^40^ to produce excitatory, inward currents that are mediated by α-MSH. To focus on melanocortin signaling, the non-selective opioid antagonist naloxone (1 μM) was present in the bath to block the potential postsynaptic effects of β-endorphin release. When a low frequency (glutamatergic) response was seen in a putative Kiss1^ARC^ neuron with a stable baseline, the slice was optogenetically stimulated at 20 Hz for 10s. This stimulation protocol has been effective for peptide release from both Kiss1 and POMC neurons ^35,36^. The current was monitored for 50 s with a brief break to run a voltage step protocol. As predicted, this stimulation paradigm frequently (35%, 7/20) resulted in a rapidly developing inward current (**Figure 6H**) that, based on the current/voltage plot, was due to the activation of a cation conductance (*data not shown*). The recording was then resumed for up to 15 m. In four cases, the inward current took several minutes to fully develop (**Figure 6I**).

Based on our findings, we hypothesized that the high frequency response was due to evoked peptide release and postsynaptic activation of melanocortin receptors. To confirm this, in a subset of the previous recordings with a high frequency response, the slice was perfused with the MCR3/4 antagonist SHU 9119 (20 nM) for >15 minutes (**Figure 7A**, right trace). This treatment reversed the current (**Figure 7B right**). Even when relatively short latency inward currents were recorded (**Figure 7C** left), SHU treatment eliminated the evoked responses (**Figure 7C** right). SHU was effective in blocking the “slow” inward currents in all cells in which it was tested (n=4) (**Figure 7D**). However, it should be noted that inward currents seen only during high frequency stimulation (directly under the blue bar) were not affected by SHU, as these were likely due to glutamatergic signaling (**Figure 7E**).

**Figure 7.**
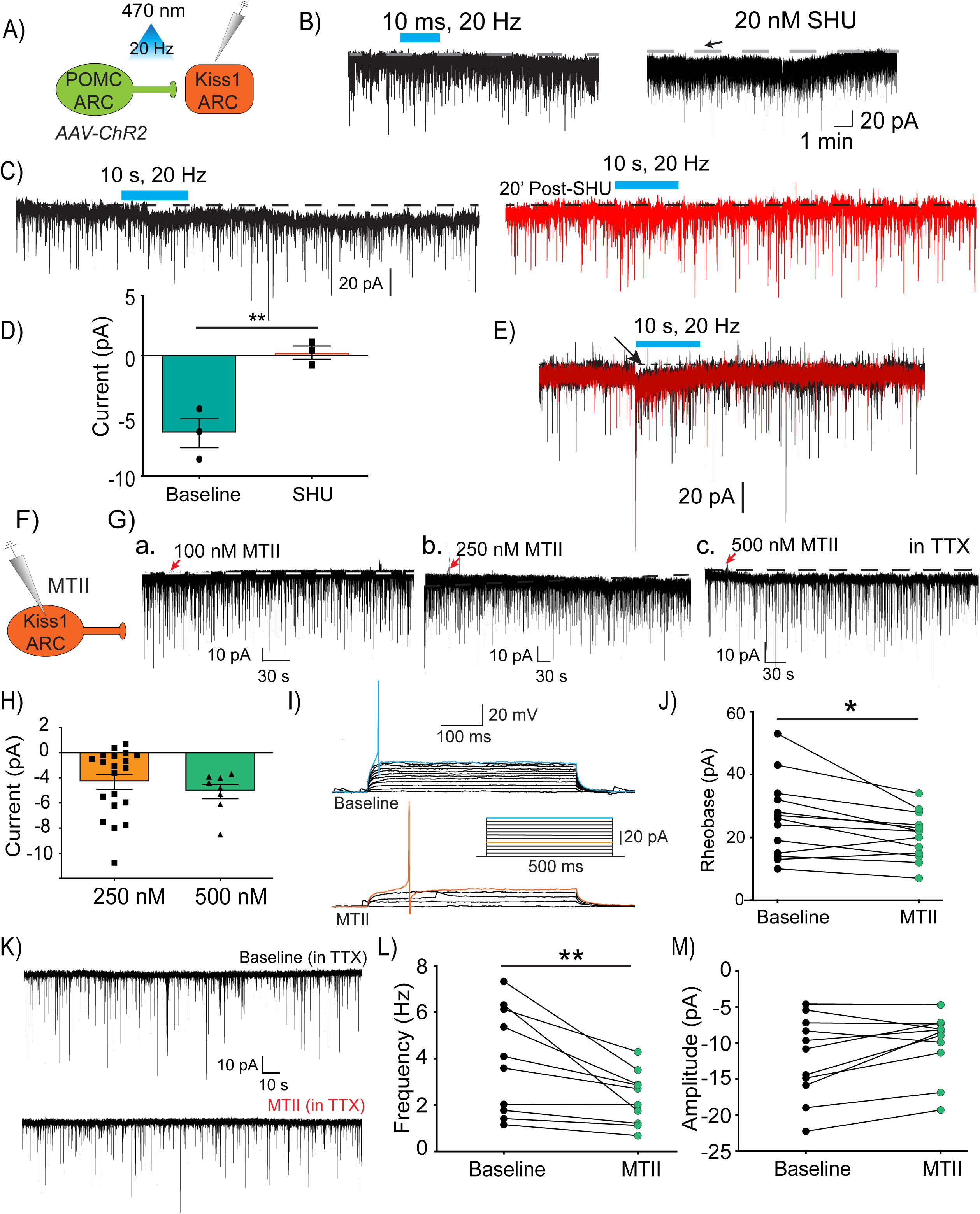
POMC neurons activate Kiss1^ARC^ neuron through direct actions on MC4R. **A)** POMC neurons were optogenetically stimulated while recording from Kiss1^ARC^ neurons. **(B)** High-frequency stimulation elicited sustained inward currents that were reversed by application of the MCR antagonist SHU. **(C)** SHU treatment also prevented subsequent stimulation of an inward current following recovery of the cell. **(D)** Bar graph comparing average response to high-frequency stimulation during baseline or following SHU treatment (Paired t-test, t=10.24, df=3, p<0.01). **(E)** The inward current present during optogenetic stimulation was not sensitive to SHU. **(F)** Recordings were made from EYFP Kiss1^ARC^ cells during application of the MCR agonist melanotan II (MTII). **(Ga)** 100 nM MTII application did not result in a clear response (1.9±1.3 pA, n=4). **(Gb-c)** 250 and 500 nM MTII were capable of eliciting similar inward currents **H)** Bar graphs comparing MTII-induced currents using 250 or 500 nM MTII (-1.8±0.8 vs -2.0±0.6, unpaired t-test, t=0.1854, df=31, p>0.05). **(I)** Representative traces in current clamp before (top) and after MTII treatment (bottom). Current was injected in incremental steps (inset) until an action potential was observed (rheobase). **(J)** Line graph showing a significant change in rheobase following MTII treatment, indicating increased excitability. Paired t-test (t=2.805, df=12, p<0.05). **(K)** Representative traces of mEPSCs in a Kiss1 neuron before and after MTII treatment. **(L)** ­Line graph showing a significant change in frequency of mEPSCs before and after MTII treatment, suggesting a decrease in spontaneous presynaptic glutamate release. Paired t-test (t=3.298, df=9, p<0.01). **(M)** Line graph of average mEPSC amplitude before and after MTII treatment, indicating no effect on postsynaptic glutamate receptor conductance or number. Paired t-test (t=2.215, df=10, p<0.05)

Having established that POMC neurons made synaptic contact with Kiss1^ARC^ neurons and often exhibited slow responses to high frequency stimulation, we wanted to confirm that activation of melanocortin receptors increases the excitability of Kiss1^ARC^ neurons. For these experiments we used Kiss1-Cre::Ai32 mice ^36^ to enable easy visualization of Kiss1 neurons (**Figure 7F**). TTX (1 μM) was maintained in the bath to block voltage-gated sodium channels, electrically isolating the neurons. Once a cell was patched with a stable membrane potential, the high affinity melanocortin receptor agonist melanotan (MTII) was directly added to the slice chamber as a microdrop (10 μl into 1 ml total volume). With 100 nM MTII (final bath concentration), there was very little effect on the membrane holding current (, **Figure 7G-a**). However, increasing the concentration up to 250 nM MTII invariably generated a slowly developing inward current (**Figure 7G-b**). Finally, 500 nM MTII produced a more rapidly developing inward current, which was most likely due to MTII’S pharmacokinetics (**Figure 7Gc**). However, there was not a significant difference in the efficacy of the two concentrations of MTII (**Figure 7H**). Given that MCR4 is Gαs-coupled ^46,47^, which robustly increases cAMP levels, we sought to determine if MTII caused a significant increase in Kiss1 neuronal excitability. Since HCN (“pacemaker”) channels are a prominent target of downstream Gαs signaling based on their cyclic nucleotide binding site ^48^, we utilized a simple measure of cell excitability—*i.e.,* rheobase, which is the minimum injected current that is required to fire an action potential. Therefore, we made current clamp recordings from Kiss1::Ai32 cells before and after microdrop application of 500 nM MTII. Current was continuously injected to keep Kiss1^ARC^ neurons around -80 mV to limit the spontaneous firing of action potentials. Next, positive current was injected in small steps until an action potential was generated, approximately 10 minutes after MTII application. We found the rheobase was significantly reduced by MTII (**Figure 7I, J)**, indicating an increased excitability likely due to elevated cAMP levels that targeted excitatory ion *(e.g.,* HCN) channels. Together these results show that melanocortin receptor activation has significant excitatory effects on Kiss1^ARC^ neurons.

Finally, voltage clamp recordings were made in the presence of TTX to measure miniature excitatory currents (mEPSCs) before and after a microdrop of MTII in the bath (10 minutes, final concentration 500 nM, **Figure 7K**). Both the frequency and amplitude of mEPSCs over at least three minutes were calculated. The frequency of mEPSCs was significantly decreased, but the average amplitude remained unchanged (**Figures 7L-M**). Changes in frequency are interpreted as reduced presynaptic release probability ^49^, whereas alterations in amplitude are thought to reflect differences in postsynaptic receptors ^50^. However, MTII decreased presynaptic glutamate release but had no effect on postsynaptic response, which would suggest that the excitatory effects of MTII do not rely on greater excitatory input from presynaptic neurons or enhanced response of postsynaptic ionotropic glutamate receptors.

## DISCUSSION

Our findings reveal a previously unrecognized melanocortin-responsive circuit that regulates energy expenditure through a subset of glutamatergic Kiss1 neurons in the arcuate nucleus. We demonstrate that these neurons are both direct targets of POMC input and integral components of a thermoregulatory pathway that projects to CART/leptin receptor sensitive neurons in the DMH and subsequently activates BAT thermogenesis via the RPa. Notably, we identify that this pathway is dependent on MC4R signaling within Kiss1^ARC^ neurons. These findings extend the current understanding of MC4R-mediated metabolic regulation, which has largely centered on distinct populations in the PVH and brainstem, by identifying Kiss1^ARC^ neurons as a node linking melanocortin input to thermogenic output ^20,21,51^.

While MC4R is well established as a central regulator of both appetite and energy expenditure ^52^, these functions are canonically assigned to distinct neuroanatomical populations: MC4R^PVH^ neurons mediate satiety, and MC4R expressed on spinal autonomic neurons facilitates thermogenesis ^21,51^. Our findings introduce Kiss1^ARC^ neurons as an additional MC4R-expressing population that is necessary for the thermogenic branch of this system. The deletion of MC4R in Kiss1 neurons resulted in impaired BAT activation, reduced *Ucp1* expression, and obesity, all occurring without changes in food intake, locomotor activity, or reproductive function (these mice presented normal LH and testosterone levels). These results delineate a cell-type-specific contribution of Kiss1 neurons to metabolic regulation, independent of their established roles in reproduction. Moreover, they reveal a sex-specific contribution of these MC4R expressing Kiss1 neurons in the regulation of energy homeostasis, as deletion of MC4R from Kiss1 neurons in female mice does not affect body weight under regular chow ^33^, although an exhaustive metabolic characterization of female Kiss1^MC4RKO^ mice remains to be performed. Additionally, Kiss1-specific MC4R null mice displayed normal glucose metabolism. This is in line with previous studies documenting that MC4R participates in the control of glucose tolerance at the level of the lateral hypothalamic neurons ^53^, which remained intact in Kiss1^MC4RKO^ mice. The loss of normal EE observed in Kiss1^MC4RKO^ mice indicate that MC4R expression in Kiss1 neurons is required for this regulatory function, establishing Kiss1 neurons as a melanocortin sensitive node in the hypothalamic control of metabolism.

In addition to altered energy expenditure, Kiss1^MC4RKO^ mice exhibited a disruption in circadian rhythmicity of feeding and drinking behaviors, characterized by a loss of the normal light-dark cycle in food and water intake. This is in line with previous findings showing that Kiss1^ARC^ neurons orchestrate the daily timing of feeding ^18^. The phenotype of our Kiss1^MC4RKO^ mice suggests that Kiss1 neurons receive a circadian input from the melanocortin system and, in turn, contribute to the circadian regulation of metabolic rhythms.

Mechanistically, our data support a model wherein Kiss1^ARC^ neurons integrate upstream melanocortinergic input from POMC neurons and transmit this signal to the DMH via glutamatergic projections. Similar to female mice ^24^, we found POMC inputs to Kiss1^ARC^ neurons lead to, not only rapid excitatory input, but greater excitability. Activation of G-protein coupled melanocortin receptors induced an inward, depolarizing current and reduced the current required to elicit an action potential. Therefore, the efficacy of other excitatory inputs to Kiss1^ARC^ neurons will also be enhanced. As with female mice ^44^, this excitatory drive is then relayed to the DMH. We then showed that the DMH neurons receiving this input—identified as leptin-sensitive and CART-expressing—project to the RPa. This is in line with previous reports demonstrating that the glutamatergic subset of Lepr^DMH^ neurons regulates thermogenesis through their connections with RPa ^27,28^, suggesting that this population is activated by Kiss1^ARC^ neurons. Activation of Kiss1^ARC^ neurons produced rapid increase in BAT temperature, while targeted inhibition of their terminals in the DMH blunted this effect, thus demonstrating a direct effect at the DMH and not through potential collaterals. Importantly, ablation of Vglut2 in Kiss1 neurons led to similar BW increases to those seen with Kiss1 neuron ablation, underscoring the requirement for glutamatergic neurotransmission in this neural circuit. This pathway is present in both sexes, as we previously reported that glutamatergic activation of Kiss1^ARC^ neurons terminals in the DMH activate Lepr/CART neurons in female mice ^44^.

Although previous work has suggested that POMC neurons drive thermogenesis, including through rapid activation of BAT ^26^, and Kiss1 neurons synapse onto POMC neurons ^54^, our findings argue that Kiss1 neurons do not mediate their metabolic effects via a Kiss1→POMC axis, but rather they convey the action of POMC neurons. First, we show that POMC neurons form functional monosynaptic inputs onto Kiss1^ARC^ neurons. Second, loss of MC4R signaling specifically in Kiss1 neurons impairs thermogenesis and leads to obesity despite intact POMC circuitry. Third, our anatomical mapping reveals robust projections and activation of Kiss1^ARC^ neurons to CART/Leptin sensitive neurons in the DMH.

The absence of changes in food intake after activation of Kiss1^ARC^ neurons align with those of Fenselau et al. ^26^, who reported that acute activation of Kiss1^ARC^ neurons does not influence food intake, reinforcing the notion that this population is selectively involved in the regulation of energy expenditure, rather than feeding behavior. Instead, we propose that Kiss1 neurons act downstream of POMC input, forming a distinct arm of the melanocortinergic system dedicated to energy expenditure.

These data also clarify the broader role of Kiss1 neurons in energy homeostasis. While prior studies have linked global kisspeptin signaling to metabolic phenotypes ^8^, we show that Kiss1^ARC^ neurons can modify energy expenditure in a kisspeptin-independent manner through glutamate release. This finding aligns with emerging literature describing non-peptidergic roles for Kiss1 neurons in hypothalamic circuitry and establishes a mechanistic link between energy balance and a classically reproductive neuronal population ^18,36^.

In summary, we identify Kiss1^ARC^ neurons as a distinct population of MC4R-expressing neurons that links melanocortin input to thermogenic output via direct glutamatergic projections to the DMH. This circuit promotes energy expenditure without affecting feeding or reproductive function, highlighting a functionally specialized role for Kiss1 neurons in the regulation of energy homeostasis.

## RESOURCES AVAILABILITY

### Lead contact

Victor M. Navarro.

### Materials availability

No unique reagents were generated in this study.

### Data and code availability

Any additional information required to reanalyze the data reported in this paper is available from the lead contact upon request.

## ACKNOWLEDGEMENTS

The authors would like to thank Dr. Brad Lowell (Beth Israel Deaconess Medical Center) for providing the MC4R floxed mouse line and Dr. Alex Banks (Beth Israel Deaconess Medical Center) for the helpful discussions and assistance with the CLAMS experiments. We also thank Isabel López for her excellent technical assistance in taking care of mouse colonies, genotyping and slicing the brains, Dr. Rona Carroll for her assistance at Harvard Medical School, and Perla Alpizar Chacon and Cecilia Cuilty Villareal for their excellent technical assistance. The University of Virginia Center for Research in Reproduction Ligand Assay and Analysis Core is supported by the Eunice Kennedy Shriver NICHD/NIH (NCTRI) Grant P50-HD28934. This work was supported by the NIH grants P30-MH048736 and R01-MH104450 to L.S.Z, the Boston Nutrition Obesity Research Center Pilot & Feasibility award # 3P30DK046200-29, The Mathers Foundation # MF-2204-02555 and Nutrition Obesity Research Center at Harvard # NIDDK-P30-DK040561 to N.L.S.M, the PHS MPI grant DK68098 to M.J.K and O.K.R; and HD090151, HD099084 and DK133760 to V.M.N. T.L.S received some support (startup funds) from Appalachian State University. R.T was supported by the Charles A. King Trust Postdoctoral Research Fellowship Award, the Lalor Foundation Postdoctoral Fellowship Award, the Women’s Brain Initiative Fellowship Award, the IBRO-ISN Research Fellowship Award, the Shore faculty career development award, and the ROSA SCORE pilot grant (supported by NIH Research Grant U54 AG062322 funded by The National Institute on Aging (NIA) and Office of Research on Women’s Health (ORWH)).

The figures and graphical abstract uses resources from the Paxinos atlas and https://scidraw.io.

## AUTHOR CONTRIBUTIONS

RT, TLS, MJK, NLSM and VMN conceived the study, designed the experiments, analyzed the data and wrote the manuscript with input from ET, KF, CJH, EM, KW, SZ, SAP, CEM, and OKR. RT, TLS, NL, ET, KF, CJH, EM, KW, SZ, SAP, CEM, NL and NLSM performed the experiments. L.Z, provided the AAV1-FLEX-Sacas9-sgSlc17a6, the AAV1-FLEX Sacas9-Sg ROSA 26, and the AAV1-FLEX KASH EGFP.

## METHODS

### EXPERIMENTAL MODEL AND STUDY PARTICIPANT DETAILS

#### Mice

*All in vivo experiments* were conducted in accordance with the guidelines of the Harvard Medical Area Standing Committee on the Use of Animals in Research and Teaching. Adult (3-8 months) males Kiss1-neuron MC4R knock-out (Kiss1^MC4RKO^) ^24^, Kiss1^Cre:GFP^ heterozygous (Kiss1^Cre/+^) and Kiss1^Cre:GFP^ homozygous (Kiss1^Cre/Cre^) mice ^55^ and their Wild-type (WT) littermates were generated. Homozygous mice (Kiss1^Cre/Cre^) are Kiss1KO and display hypogonadotropic hypogonadism ^55^. Unless otherwise stated, mice were group housed in Harvard Medical School Animal Resources facilities under constant conditions of temperature (22–24°C) and light (12:12 h light-dark cycle). Mice had *ad libitum* access to water and either regular chow (*Teklad F6 Rodent Diet 8664*) or 60% high-fat diet (HFD, 60% fat. *Fisher Scientific, Cat. NC0004611*). Littermates of the same sex were randomly assigned to experimental groups.

*All in vitro experiments* were performed in accordance with institutional guidelines based on National Institutes of Health standards and approved by the Institutional Animal Care and Use Committees at Oregon Health and Science University and Appalachian State University. Kiss1^Cre:GFP^ mice ^55^ were crossed with Ai32 ^56^ or C57B6J mice. POMC^Cre^ mice (RRID: JAX:005965) ^57^ were crossed with wildtype C57B6J (RRID: JAX:000664) mice. The Ai32 cross was not used with POMC^Cre^ mice because the gene is transiently expressed during development in some cells fated to be Kiss1 or AgRP cells ^58^ and early recombinant events lead to persistent expression of the ChR2-mCh fusion protein in non-POMC cells. However, we have previously shown that AAV-driven expression in adult POMC^Cre^ animals is restricted to b-endorphin labeled cells in the ARC^35^, avoiding this specificity issue. All colonies were maintained onsite under controlled temperature (21-23 °C) and photoperiod (12:12-h light-dark cycle 0600 to 1800) while receiving ad libitum food (5L0D; LabDiet, St. Louis, MO) and water access. Following surgeries, mice received a *s.c.* dose of 4-5 mg/kg carprofen (Rimadyl; Pfizer Animal Health, New York, NY) and given at least 1 week of recovery.

#### Stereotaxic surgeries

Mice were deeply anesthetized with isoflurane and placed into a stereotaxic apparatus (Kopf Instruments, Model 940). For postoperative care, mice were injected subcutaneously with sustained-release meloxicam (5mg/kg) and buprenorphine (0.6 mg/kg). Lubrication was applied to the eyes, and the surgical site was cleaned with betadine and ethanol. After exposing the skull via small incision, a small hole was drilled for injection at the appropriate anterior-posterior (AP) and medio-lateral (ML) coordinates. A syringe (Hamilton, 2.0 µL, Neuros Syringe, Model 7002 KH, Ref. PN: 65459-01) was lowered into the brain at the appropriate dorsal-ventral (DV) coordinates. Injection sites were chosen based on the Paxinos Brain Atlas, and confirmed with India Ink (Fisher Scientific, Cat. NC9903975) in trial injections. Each infusion was slowly delivered (100 nl/min), and the needle was left in place for an additional 5 min and then slowly withdrawn to minimize backflow.

***For Kiss1^ARC^ neurons ablation experiments***, AAV5-flex-taCasp3-TEVp (Addgene, Cat. 45580, 1×10^13^ GC/mL) was injected bilaterally in adult Kiss1^Cre/+^ mice in the ARC (co-ordinates from bregma: AP= -1.6 mm, ML= ±0.25 mm, DV= -5.85 mm; 200nl per site). A control group of Kiss1^Cre/+^ mice was injected with AAV5-hSyn-DIO-mCherry (Addgene, Cat. 50459. 8.4×10^12^ GC/mL). Mice were allowed to recover in their home cages and body weights were measured twice a week for 12 weeks.

***For Kiss1^ARC^ chemogenetic experiments***, AAV9-hSyn-DIO-hM3D(Gq)-mCherry (Addgene, Cat. 44361, 2.6×10^13^ GC/mL) was injected bilaterally in Kiss1^Cre/+^ adult mice in the ARC (coordinates and volume as above). A control group of Kiss1^Cre/+^ mice was injected with the AAV5-hSyn-DIO-mCherry. Mice were allowed recovery and AAV expression for at least 3 weeks and chemogenetic activation of Kiss1^AR^ neurons was achieved with intraperitoneal injection of Clozapine-N-Oxide (CNO, 3mg/kg).

***For RPa retrograde tracing experiments***, Green Retrobeads™ IX (Lumafluor, 50 nl) was injected in the RPa (coordinates: AP= -6 mm, ML= 0.0, DV= -5.8 mm from brain surface) in Kiss1^Cre/+^ adult mice. 5 days following injections animals were perfused.

***For CRISPR Mutagenesis of Vglut2 in Kiss1^ARC^ Neurons***, CRISPR/SaCas9 viruses were prepared at the University of Washington as previously described ^34^. While incorporating the SaCas9 sequence allowed the inclusion of the sgRNA in a single vector, there was not sufficient space remaining in the cassette for a fluorophore sequence. Therefore, to visualize the quality and location of the injection/infection, 90% of the AAV1-FLEX-Sacas9-sgSlc17a6 (University of Washington) was spiked with 10% of a high-titer virus encoding EGFP (AAV1-FLEX KASH EGFP, University of Washington). We have previously validated and shown that this mixture is able to successfully infect Kiss1 neurons in Kiss1^Cre/+^ mice and significantly decrease Vglut2 expression in Kiss1 ARC neurons ^33^. The CRISPR/SaCas9 mixture was bilaterally injected in the ARC (coordinates and volume as above) of adult (3 month) Kiss1^Cre/+^ mice. Control Kiss1^Cre/+^ mice were bilaterally injected with 90% AAV1-FLEX Sacas9-Sg ROSA 26 (University of Washington, titer xxx) spiked with 10% AAV1-FLEX KASH EGFP. Mice were allowed to recover in their home cages and body weights were measured twice a week, first under regular chow (for 12 weeks), then under high fat diet (for 8 weeks).

***For in vivo optogenetic inhibition of Kiss1 fibers in the DMH****, AAV1-hSyn1-SIO-eOPN3-mScarlet-WPRE* (Addgene, Cat. 125713, 2.2×10^13^ GC/mL) *was injected bilaterally in the ARC* (coordinates and volume as above) of Kiss1^Cre/+^ mice. This AAV causes Cre-dependent expression of the eOPN3, an inhibitory opsin that permits long-term inhibition without damaging the tissue. For optogenetic manipulation of Kiss1^ARC^ terminals, an optic fiber (200 µm diameter core; 0.50NA, Ø1.25 mm Ceramic Ferrule. Model: R-FOC-L200C-50NA, RWD Life Science Inc) was implanted unilaterally over the DMH in the middle line (above the third ventricle) at the coordinates: AP= 1.85 mm, ML= 0.0, DV =4.45 mm from Bregma. The experimental group consisted of animals injected with the AAV-eOPN3 and exposed to photoinhibition (INHIB), while the control group consisted of the same mice but without light delivery (SHAM).

***For the mapping of Kiss1^ARC^ neurons axonal projections***, we used the brains injected with AAV1-hSyn1-SIO-eOPN3-mScarlet-WPRE in our optogenetic experiments. This vector presents robust and specific expression in Kiss1^Cre^ neurons and strong labeling of long-range axonal projections.

***For in vitro optogenetic stimulation of Kiss1 fibers in the DMH***, bilateral ARC injections of AAV1-Ef1a-DIO-ChR2: mCherry (Addgene, Cat. 20297, 2.0×10^12^ particles/ml; 400nl per site) or AAV1-Ef1a-DIO-ChR2: YFP (Addgene, Cat. 202 98, 2.0x10^13^; 400nl per site) were performed on adult Kiss1^Cre^ or POMC^Cre^ male mice. ARC injection coordinates were: AP= −1.10 mm, ML= ± 0.30 mm, DL=−5.80 mm (from surface of brain z = 0.0 mm). Mice were given carprofen for analgesia and allowed to recover for at least two weeks before euthanasia and tissue harvesting.

### In vitro electrophysiology

#### Visualized whole-cell patch recordings

Hypothalamic coronal brain slices (240 µm) were made from male mice, using a Leica VT1000S vibratome, in ice cold cutting solution bubbled with O_2_/C0_2_ (95%/5%). Slices were then transferred to a holding chamber with artificial cerebrospinal fluid bubbled with the same gas mix and allowed to recover for at least 1 hour. For recordings, slices were placed in a perfusion chamber and visualized with an Olympus BX51W1 using either differential infrared contrast or oblique illumination. Electrodes were fabricated with a Sutter Instruments Puller (P-97 or P-1000) from borosilicate glass (1.5-mm outer diameter, World Precision Instruments). Pipette resistances ranged from 3-5 MΩ and access resistance was less than 20 MW. Electrophysiological signals were amplified with an Axopatch 200A/B amplifier and digitized with a Digidata 1440A/1550B (Molecular Devices) using pClamp (10/11; Molecular Devices). The liquid junction potential was corrected for all recordings.

Kiss1 neurons in the ARC were targeted based on fluorescence (Ai32 cross) or as done previously and described below ^29,59^ for POMC^Cre^-injected animals. Final concentration was calculated based on the drug in the known volume of the bath. Perfused drugs (TTX, SHU9199, and naloxone) were constantly circulated and given at least 10 minutes to reach maximal effect. Focal application of drugs (MTII) was done with the pump stopped (10-20 minutes). 0.3 to 1 ml was directly added to the known volume of the bath to achieve the desired final concentration. This approach enables precise timing of recordings and reduces the risk of receptor desensitization. For optogenetic stimulation, a light-induced response was evoked using a 470 nm LED and driver (ThorLabs) with the light path directly delivered through the 40X water-immersion lens.

#### Targeting of Kiss1 neurons for electrophysiological recordings

For non-optogenetic experiments, brain slices were taken from AAV injected Kiss1^Cre^ or Kiss1^Cre^xAi32 male mice. Ai32 mice (RRID:IMSR_JAX:024109, C57BL/6 background) carry the floxed ChR2 (H134R)-EYFP gene in their Gt(ROSA)26Sor Locus ^56^, allowing its expression in a Cre-dependent manner. Due to concerns of nonspecific expression ^60^, we previously validated this model using single cell RT-PCR and documented that *Kiss1* mRNA was detectable in 99% of individually harvested eYFP cells (n=126). In addition, we have used both AAV injected Kiss1-Cre AAV or Kiss1xAi32 male mice and found no differences in electrophysiological results ^59^. We used AAV injected POMC^Cre^ male mice and avoided small soma, high input resistance (>800 MΩ), ventrally located cells that are typically NPY/AgRP neurons. Instead, we targeted larger, more dorsomedial neurons in the ARC while avoiding fluorescent (POMC) cells. Unlike ARC Kiss1 neurons, POMC neurons do not make reciprocal projections, so uninfected POMC neurons do not display monosynaptic EPSCs in response to optogenetic stimulation. Next, we used a ramp IV protocol to probe for the presence of a persistent sodium current (I_NaP_). While this current is more prevalent in the AVPV Kiss1 population, Kiss1 neurons are the only ARC neurons to display this electrophysiological “fingerprint” of a pronounced I_NaP_ paired with a high capacitance and low input resistance ^45^. Finally, we also harvested the cytosol at the end of all recordings and measured the expression of Kiss1. Only cells that expressed a persistent sodium current and/or expressed *Kiss1* mRNA were included in final analysis.

#### Electrophysiology data analysis

Electrophysiological data were analyzed using Clampfit 10/11 (Molecular Devices), MiniAnalysis (Synaptosoft), and Prism 7/10 (Dotmatics). All values are expressed as Mean ± SEM. Comparisons between two groups were made using un-paired Student’s t-test or between multiple groups using an ANOVA (with *post hoc* comparisons) with p-values < 0.05 considered significant. When variances differed significantly, Mann-Whitney U test was used instead.

#### Solutions/drugs

Standard vibratome slicing, external, and internal recording solutions were utilized as previously described ^29,59^. Tetrodotoxin was purchased from Alomone Labs (Jerusalem, Israel), Melanotan II, αMSH, SHU9199, and CNQX from Tocris (Minneapolis, MN). TEA, 4-AP, and Naloxone were purchased from Millipore-Sigma.

### iBAT temperature

***For the chemogenetic activation of Kiss1^ARC^ neurons***, adult Kiss1^Cre/+^ and Kiss1^Cre/Cre^ mice were injected with either AAV9-hSyn-DIO-hM3D(Gq)-mCherry or AAV5-hSyn-DIO-mCherry (for control groups). Two weeks later, they were implanted with Star-Oddi DST (data storage tag) Nano-T temperature probe (EMKA Technologies, Falls Church, VA) subcutaneously above the iBAT pad between the scapula under anesthesia. Animals were allowed 1 week recovery and acclimation was performed with intraperitoneal injection of saline for 7 consecutive days. Mice were individually housed in their home cages with free movement and *ad libitum* access to food and water. During experiments, iBAT temperature was recorded every 5 min following CNO injections (3mg/kg). The temperature was captured by the temperature logger implanted in each animal and was extracted using the Mercury software. BAT temperature data were analyzed as ΔT (delta temperature), calculated as the change in temperature from baseline (time 0), defined as the time of CNO injections. For each mouse, baseline temperature at time 0 was subtracted from subsequent temperature values to assess the effect of CNO over time. Group average of ΔT were then computed and compared across experimental conditions.

### Thermal imaging

***For the optogenetic inhibition of Kiss1^ARC^ fibers in the DMH*,** thermal imaging was performed in Kiss1^Cre/+^ mice using an infrared camera (FLIR E4). Before the recordings, the back, neck, and head of the mouse were shaved under a brief exposure to isoflurane. Following at least 2 days of recovery from anesthesia and 24 hours for habituation, photoinhibition was performed. We recorded the temperature for 30 minutes baseline, 1 hour photoinhibition (INHIB), and 1 hour recovery. The analysis was performed every 15 minutes. All recordings were performed during the light phase and the animals were not disturbed prior to the onset of the experimental protocol. Thermal images from videos recorded the time course changes in iBAT temperature and were used to determine ΔT iBAT using a mean of the temperature between the right and left scapula from each mouse. Group average of ΔT were then compared. FLIR tools software (V6.4, 2015) was used for all analysis.

### Photoinhibition

Photoinhibition was delivered by 10 ms pulses at 4–8 mV. The blue light laser (470 nm; LaserGlow) was controlled by Spike 8.01 software using a 10 Hz stimulation pattern of 2 sec-on and 3-off, as described in ^32,61^, which causes no evidence of injury to the target neurons.

### In situ hybridization

For a maximal expression of *Kiss1* mRNA in the ARC, WT and Kiss1^MC4RKO^ male mice were castrated and sacrificed one week later by decapitation. Brains were extracted fresh frozen on dry ice and stored at -80°C until sectioned. Five sets of 20 µm of coronal sections were cut on a cryostat, from the diagonal band of Broca to the mammillary bodies, mounted onto SuperFrost Plus slides (VWR Scientific) and stored at -80°C until use. A single set was used for the experiments (adjacent sections 100 mm apart). Dual fluorescence *in situ* hybridization was performed using RNAscope (ACD bio, Multiplex Fluorescent v.2) according to the manufacturer’s protocol using the following probes: *Mc4r* (319181-C2) and *Kiss1* (500141-C1). Fluorescent images were taken using a Zeiss Axio Imager M2 Microscope.

### Histology

#### Brain tissue preparation

Mice were euthanized by deep anesthesia with intraperitoneal injection of either a ketamine/xylazine cocktail in saline (0.9%) or a chloral hydrate (1.5% BW, 7% solution). They were transcardially perfused with 30ml of 0.1M phosphate buffer (0.1M PB) followed by 30ml of 4% paraformaldehyde diluted in 0.1M PB (PFA; Boston BioProducts). Brains were extracted and post fixed overnight in PFA and then stored in 20% sucrose (Thermo Fisher Scientific) until sectioned, using a freezing microtome (30µm coronal sections into 4 series). Following sectioning, tissue was stored at -20°C in a cryoprotectant solution [30% sucrose in 0.1M PB containing 30% ethylene glycol (Thermo Fisher Scientific) and 0.01% sodium azide] until further processing.

#### mCherry and cfos staining

To assess the expression of the mCherry reporter in Kiss1^ARC^ neurons and their projections, and to analyze cfos expression in the DMH, AAV9-hSyn-DIO-hM3D(Gq)-mCherry or AAV5-hSyn-DIO-mCherry mice were injected with CNO (3mg/kg) 90 min before perfusion for cfos visualization and quantification. Free-floating coronal sections were incubated with rat anti-mCherry conjugated to Alexa Fluor 594 (Thermo Fisher, Cat. M11240; 1:500) and guinea pig anti-cfos (Synaptic Systems, Cat. 226 308; 1:1000) for 48 hours at 4°C and for 2 hours with donkey-anti-guinea pig Alexa 488 (Jackson ImmunoResearch, Cat. 706-545-148; 1:400). Sections were washed, mounted on microscope slides air-dried and cover slipped with Vectashield Mounting Medium (Vector Laboratories, Burlingame, CA). Total number of cfos positive neurons in the DMH of Kiss1^Cre/+^ and Kiss1^Cre/Cre^ mice was quantified manually in images taken at x20 magnification from DMH sections between bregma -1.70 mm and -2.60 mm (1 section/animal).

#### mCherry, cfos and pStat3 staining

To investigate whether Kiss1^ARC^ neurons projections contact leptin receptor expressing neurons in the DMH and induce cfos expression in these neurons, a tiple mCherry/cfos/pStat3 immunohistochemistry was carried out. Kiss1^Cre/+^ mice were injected with AAV9-hSyn-DIO-hM3D(Gq)-mCherry, and 3 weeks later mice were fasted overnight. The next day, mice were injected with i) CNO (3mg/kg) 90 minutes before perfusion (to analyze cfos expression), and ii) recombinant murine leptin (PeproTech, Cat. 450-31; 5 mg/kg) 30 minutes before perfusion (to induce phospho-Stat3 (pStat3) expression). Mice were perfused with PBS followed by 4% PFA and sections were post-fixed, cryoprotected in 30% sucrose and sectioned at 30 µm, as described above. Free-floating sections underwent antigen retrieval in10mM sodium citrate buffer (PH=6) in 90°C for 20 min to enhance pStat3 signal. After cooling, sections were blocked in PBS [containing 0.3% Triton, 0.1% bovine serum albumin (BSA), 5% normal goat serum (NGS) and 5% normal donkey serum (NDS)] and were incubated sequentially with the following primary antibodies: guinea pig anti-cfos (Synaptic Systems, Cat. 226 308; 1:1000), rat anti-mCherry conjugated to Alexa Fluor 594 (Thermo Fisher, Cat. M11240; 1:500) and rabbit anti-pStat3 (Cell Signaling Technology, Cat. 9145L; 1:250) for 48 hours at 4°C. Section were then incubated with the secondary antibodies: Alexa Fluor 647 donkey-anti-guinea pig (Jackson ImmunoResearch, Cat. 706-605-148; 1:200) and DyLight 488 goat-anti-rabbit (Thermo Scientific, Cat. 35552; 1:200). Confocal images were taken at the Havard Medical School Imaging Core Using the Leica Stellaris X5.

#### GFP fluorescent staining

To validate the targeting and infection of Kiss1^ARC^ neurons in the CRISPR-Cas9 mutagenesis experiments, we performed immunofluorescent staining for GFP. As mentioned above, the SaCas9 sequence allowed inclusion of the sgRNA in a single vector but left insufficient space for a fluorescent reporter. To allow histological visualization of infection, a high titer AAV encoding GFP (AAV1-FLEX KASH EGFP) was co-injected with the SACas9 virus. Free-floating coronal sections were stained with a rabbit-anti-GFP (Thermo Fisher, Cat. A-6455; 1:5000) for 48 hours at 4°C, followed by goat-anti-rabbit DyLight 488 secondary antibody (Thermo Scientific, Cat. 35552; 1:200).

#### RFP staining

To validate the targeting and infection of Kiss1 neurons with the AAV1-hSyn1-SIO-eOPN3-mScarlet-WPRE, we performed immunofluorescent staining against the red fluorescent protein (RFP) as the inhibitory opsin eOPN3 is fused to the red fluorescent protein mScarlet. Free-floating coronal sections were incubated with a rabbit-anti-RFP (Rockland, Cat. 600-401-379; 1:500) for 48 hours at 4°C, followed by donkey anti-rabbit Alexa Fluor (Abcam, Cat. 175470; 1:500). Images were collected using the V200 Olympus slide scanner and Olyvia viewer software V4.1.

#### GFP chromogenic staining

To validate Kiss1 neurons targeting with the AAV5-flex-taCasp3-TEVp and quantify the extent of ablation of Kiss1^ARC^ neurons, we performed chromogenic immunohistochemistry against GFP. Kiss1 neurons were genetically labeled with GFP in Kiss1^Cre:GFP^ mice ^55^. Therefore, immunostaining for GFP was performed to enable visualization. Free-floating sections were incubated with rabbit-anti-GFP (Invitrogen, Cat. A-6455; 1:5000) for 48 hours at 4°C, followed by incubation in biotinylated goat-anti-rabbit antibody (Vector laboratories, Cat. BA-1000; 1:500) for 1 hour, and elite Avidin-Biotin complex (Vector laboratories, Cat. PK-6100; 1:1000). DAB (3,3′-Diaminobenzidine tetrahydrochloride, Sigma Aldrich, Cat. D5905) was used as the chromogen. For every 10mg DAB was added: 50ml 0.1M PB, 2% nickel sulfate hexahydrate (Sigma-Aldrich, Cat. 227676-100G), and 20 µl (of 30%) H_2_O_2_. Sections were incubated for 10 min in the mixture at room temperature, and were rinsed with 0.1M PB, mounted on slides and coverslipped. ARC GFP positive (Kiss1^ARC^) neurons were visualized under brightfield microscopy and were manually counted to assess the extent of Kiss1 neuronal ablation relative to control animals. All animals with unchanged number of Kiss1^ARC^ neurons were considered missed injections and were excluded from the study.

### Glucose Tolerance Test (GTT)

To assess glucose homeostasis, adult Kiss1^MC4RKO^ males and their WT littermates were fasted for 5 hours prior to the test. Mice were then injected intraperitoneally with D-glucose (Sigma-Aldrich, 2g/kg body weight) dissolved in sterile saline. Blood glucose levels were measured from tail vein blood at baseline (time 0) and at 20-, 60- and 120-minutes post-injections using a glucometer.

### Insulin Tolerance Test (ITT)

For insulin sensitivity assessment, Kiss1^MC4RKO^ males and their WT littermates were fasted for 5 hours, followed by an intraperitoneal injection of human insulin (0.75 U/kg body weight, prepared at 0.1 U/ml in sterile saline). Blood glucose levels were measured from tail bleeds at baseline (time 0) and at 20-, 40-, 60- and 120-minutes post-injections using a glucometer.

### Real time quantitative PCR

Tissues from i) BAT (for *ucp1* gene expression) and ii) preoptic area (POA), mediobasal hypothalamus (MBH), testis, pituitary gland, BAT, white adipose tissue (WAT), liver and muscle (for *Mc4r* gene expression) were used. Total RNA was extracted from frozen tissues of Kiss1^MC4RKO^ mice and their WT littermates using TRIzol reagent (Invitrogen) followed by chloroform/isopropanol extraction. RNA was quantified using a NanoDrop 2000 spectrophotometer (Thermo Scientific Scientific), and 1µg of RNA was reverse transcribed using an iScript cDNA synthesis kit (Bio-Rad). Semi-quantitative real-time PCR assays were performed on an ABI Prism 7000 sequence detection system and analyzed using ABI Prism 7000 SDS software (Applied Biosystems). The cycling conditions were as follows: 2 min incubation at 95°C (hot start), 45 amplification cycles (95°C for 30 s, 60°C for 30 s, and 45 s at 75°C, with fluorescence detection at the end of each cycle), followed by melting curve of the amplified products obtained by ramped increase of the temperature from 55°C to 95°C to confirm the presence of single amplification product per reaction. For data analysis, we used the 2(-Delta Delta C(T)) method and target genes were standardized to *Hprt* levels in each sample. The primers used are listed in Table 1.

**Table 1.**
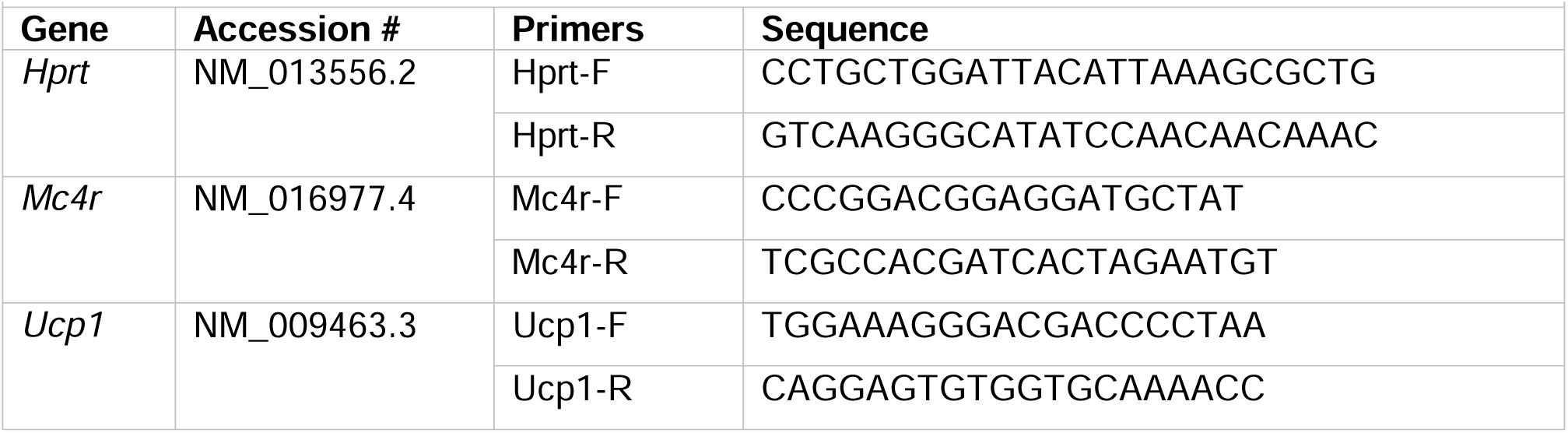
Primers Used for Polymerase Chain Reaction Assays.

### Single Cell PCR

Cells were harvested following whole-cell patch recordings and expelled into a low adhesion microcentrifuge tube containing 5X First-strand buffer (Invitrogen), 15 U Rnasin (Promega), and 100 mM dithiothreitol (DTT) in a total of 5 µl. Cells were frozen at -80°C. Cells were thawed and complementary DNA (cDNA) synthesis was performed by adding 10 mM dNTP, 100 ng random hexomers (Promega), 400 ng oligo dT (Invitrogen) and denatured for 5 minutes at 65°C, then cooled on ice for 5 minutes. Additional 5X First-strand Buffer, 15 U Rnasin and 100 mM DTT was added and 100 U of Superscript III reverse transcriptase (RT) (Invitrogen) to a final volume of 20 µl. The reaction was incubated for 5 minutes at 25°C, 60 minutes at 50°C, 15 minutes at 70°C then cooled to 4°C for 5 minutes. cDNA was stored at -20°C until PCR was performed. Control hypothalamic tissue was also processed alongside the recorded cells with and without RT. Primers for Kiss1 and CART were designed to cross at least one intron-exon boundary and are as follows: Kiss1 (120 bp product) forward primer 5’TGCTGCTTCTCCTCTGT3’, reverse primer 5’ACCGCGATTCCTTTTCC3’; CART (157 bp product) forward primer 5’AACGCATTCCGATCTACG3’, reverse primer 5’TCCCTTCACAAGCACTTC3’. Polymerase chain reaction (PCR) was performed on 3 µl of cDNA for Kiss1 and CART as follows: denature 94°C 2 minutes, 50 cycles of: 94°C for 20 s; 57°C (Kiss1), 59°C (CART) for 30 s; 72°C for 30 s; final extension at 72°C for 5 minutes. The PCR product was visualized with ethidium bromide on a 2% agarose gel.

### Oil Red O staining

Livers were collected from Kiss1^MC4RKO^ and WT littermates under regular chow. The tissues were sectioned and stained by the Rodent Histopathology Core at Harvard Medical School. Frozen sections were used for Oil Red O staining and were imaged using brightfield microscope. Lipid accumulation was quantified using ImageJ by applying a threshold and calculating the Oil Red O-positive area as previously described ^62^. % area and total lipid area per section were extracted and compared between genotypes.

### Metabolic measurements

*To assess the role of Kiss1^ARC^ neurons in the regulation of feeding behavior* **(Figure 1I)**, Kiss1^Cre/+^ males were injected with either AAV9-hSyn-DIO-hM3D(Gq)-mCherry or AAV5-hSyn-DIO-mCherry. Acute activation of Kiss1^ARC^ neurons would provide the necessary time to increase testosterone levels, however, because testosterone can significantly influence food intake, we normalized circulating testosterone levels in both groups. Two weeks following stereotaxic injections, animals underwent bilateral castration and were implanted with capsules (1.5 cm long, 0.078 in inner diameter, 0.125 in outer diameter; Dow Corning) containing testosterone in powder form (1 cm filled area). Animals were sent to the metabolic core at Beth Israel Deaconess Medical Center (BIDMC) to measure food intake at thermoneutrality (30°C). Animals were allowed an acclimation period of 24 hours in the metabolic cages under thermoneutrality and were receiving intraperitoneal injections of saline for two days before the experiment. On the experiment day, food intake was measured 4 hours before injections (to establish a baseline), CNO (3mg/kg) was injected, and food intake was recorded for another 6 hours post-injections. The experiment was performed during the light phase (6am to 6pm). The data are represented as the trace for cumulative food intake across 4 hours (figure 1, I), and the Δ FI (delta food intake), calculated as the change in food intake from baseline (time 0), defined as the time of CNO injections. For each mouse, baseline food intake at time 0 was subtracted from subsequent food intake values to assess the effect of CNO over time. Group average of Δ. FI were then computed and compared across experimental conditions.

To analyze whole body composition of Kiss1^MC4RKO^ mice and their WT littermates under regular chow **(Figure 4, E/F/G)**, adult males were euthanized using CO2 and were sent to the BIDMC metabolic core for an EchoMRI body composition analysis. % of Fat and lean mass were measured.

To investigate the origin of the obesity phenotype of the Kiss1^MC4RKO^ mice **(Figure 5)**, adult (3 months old) KO mice and their WT littermates were submitted to HFD and were sent to the Animal Physiology Core at the Joselin Diabetes Center (at 10 months old) for a Comprehensive Laboratory Animal Monitoring System (CLAMS) study. Animals were acclimatized to the metabolic cages for 48 hours, and oxygen consumption (VO2), carbon dioxide (CO2) production, respiratory exchange ratio (RER), food and drinking behaviors, activity level, and heat production were measured in the dark and light phases. Food and water intake data were analyzed as the cumulative consumption over 24 hours, while all the other parameters were analyzed as the area under the curve (AUC) to show the physiological changes over 24 hours.

### Luteinizing hormone (LH) and testosterone assays

Blood samples for LH measurements were obtained from Kiss1^MC4RKO^ males and their littermates. Briefly, the tail tip was cleaned with saline and then massaged prior to taking a 4 µl of blood sample with a pipette. Whole blood was immediately diluted in 116 µl of 0.05% PBST [phosphate buffer saline (Boston Bio Products, Cat. No. BM220) containing Tween-20 (Sigma, Cat. No. P2287)], vortexed, and frozen on dry ice. Samples were stored at -80°C until analyzed with LH ELISA as previously reported ^63^. Serum samples were also collected for the analysis of testosterone levels in adult Kiss1^MC4RKO^ males. Testosterone levels were measured at the University of Virginia Ligand Assay core with the Mouse & Rat Testosterone ELISA assay (reportable average range 10-1600 ng/dL; sensitivity of 10 ng/dL).

### Reproductive maturation of Kiss1^MC4RKO^ males

To assess their reproductive phenotype, Kiss1^MC4RKO^ mice and their control littermates were weaned at post-natal day (PND) 21 and were monitored daily for puberty onset, through monitoring of preputial separation as an indirect marker of puberty onset in the male mouse. Body weights (BW) were measured at the day of puberty onset and throughout the duration of the study.

### Castration

When necessary, 7-10 days before the experiment, gonadectomies were performed on male mice through bilateral incisions in the scrotum while under isoflurane inhalation anesthesia. The vasculature to each testicle was clamped with a small hemostat and sutured using nonabsorbable SOFSILK (MedRep Express). After the removal of each gonad, the skin incision was closed using NYLON nonabsorbable suture (MedRep Express). Each mouse received analgesia (Rimadyl; 4 mg/kg, sc) on the day of operation.

## STATISTICAL ANALYSIS

Statistical analyses were performed using GraphPad Prism v7 (GraphPad Software) and are described in the figure legends. The statistical analysis of values was performed blind to experimental conditions with statistical significance defined as P < 0.05 of values within 2 standard deviations from the mean. Size of experimental groups was determined based on historical data from our group to achieve statistical significance. Mice were randomized into control or treatment groups. Control mice were age-matched littermate where possible. All experimental animals were sacrificed after completion of the study and viral expression, injection sites, and optical fiber placement were verified. Animals with missed injections or misplacement of the optic fiber were excluded. N values are reported in the figure legends and reflect the final number of validated animals per group. Data are reported as mean ± SEM.

