## Supplemental Figures legend for "Melanocortin-responsive Kiss1 neurons of the arcuate nucleus drive energy expenditure through glutamatergic signaling to the dorsomedial hypothalamus"

**Figure S1. Anterograde Mapping of Kiss1 neurons projections**

Whole brain mapping of Kiss1 neurons projections. 3V: 3^rd^ ventricle. LV: lateral ventricle. aca: anterior commissure. f: fornix. Suprachiasmatic nucleus (SCN). +: Sparse innervation, ++: Sparse to moderate innervation, +++: Moderate-dense innervation, ++++: Dense innervation.

| **Brain region** | **Innervation** |
| --- | --- |
| Medial septum (MS) | + |
| Median preoptic nucleus (MnPO) | +++ |
| Ventro-medial preoptic nucleus (VMPO) | +++ |
| Organum Vasculosum of the Lamina Terminalis (OVLT) | + |
| Median division of the Bed Nucleus of the Stria Terminalis (BnST, M) | +++ |
| Lateral division of the Bed Nucleus of the Stria Terminalis (BnST, l) | + |
| Septohypothalamic nucleus (SHN) | +++ |
| Lateral septal nucleus (LSV) | ++++ |
| Periventricular nucleus (Pe) | ++++ |
| Anteroventral periventricular nucleus (AVPe) | +++ |
| Medial preotic area (MPA) | +++ |
| Paraventricular thalamic nucleus (PVT) | ++ |
| Paraventricular hypothalamic nucleus (PVH) | ++++ |
| Subparaventricular zone of the hypothalamus (SPZ) | +++ |
| Lateral nucleus of the hypothalamus (LH) | ++ |
| Dorso-medial nucleus of the hypothalamus (DMH) | +++ |
| Arcuate nucleus of the hypothalamus (ARC) | ++++ |
| Ventro-medial nucleus of the hypothalamus (VMH) | + |
| Postero-dorsal nucleus of the medial amygdala (MePD) | ++ |
| Postero-ventral nucleus of the medial amygdala (MePV). | + |

**Figure S2. Validation of the Kiss1^MC4RKO^ mouse**

Representative photomicrographs depicting comparable *Mc4r* mRNA expression in the PVH between WT **(A)** and Kiss1^MC4RKO^ males **(B, a)**. Scale bar is 50 µm. **(B, b)** Representative images depicting co-expression of *Kiss1* and *Mc4r* mRNA in the ARC, using RNAscope, in GDX Kiss1^MC4RKO^ male mouse. As expected, *Mc4r* was not detected within Kiss1 neurons in Kiss1^MC4RKO^ males (n=4). Scale bar is 50 µm.

**Figure S3. Reproductive phenotyping of the Kiss1^MC4RKO^ male mice**

**(A)** Cumulative percentage of puberty onset (n=13/group), and **(B)** mean age of puberty onset in in Kiss1^MC4RKO^ mice (25.55 ± 0.545, n=22) and their WT littermates (25.62 ± 0.780, n=13) show a normal puberty onset in Kiss1^MC4RKO^ males comparable to their WT littermates (p=0.470). **(C)** Kiss1^MC4RKO^ mice (12.90 ± 0.500, n=14) also show a normal body weight (BW) compared to controls (13.56 ± 0.534, n=7) at the time of puberty onset (PO), p=0.42. **(D)** Kiss1^MC4RKO^ mice (59.52 ± 19.81, n=5) also show normal testosterone levels (p=0.836) compared to their controls (66.18 ± 21.40, n=8). **(E)** Basal luteinizing hormone (LH) levels are also comparable between Kiss1^MC4RKO^ mice (0.403 ± 0.157) and WT mice (0.289 ± 0.080). n=8/group. p=0.531. Student’s t test. Data are presented as mean ± SEM.

**Figure S4. Validation of intact Mc4r gene expression in peripheral tissues**

*Mc4r* mRNA expression was similar between Kiss1^MC4RKO^ mice and their WT control littermates in the preoptic area (POA, p>0.999), the medio-basal hypothalamus (MBH, p>0.999), testis (p=0.990) and the pituitary gland (p=0.992), while it was undetected in the BAT, the white adipose tissue (WAT), the liver and muscle. Two-way ANOVA followed by Sidak’s multiple comparisons test: n=5/group.

**Figure S5. *Posthoc* identification of recorded cells**

The cytosol was harvested at the end of electrophysiological recordings and the contents underwent reverse-transcriptase and PCR with primers used to identify cell type. A) Uncropped image of gel from Figure 2N that shows cells displaying optogenetically-driven postsynaptic currents in DMH cells expressed *Cartpt*. B) Uncropped gel image from Figure 6G which shows that cells displaying optogenetically-driven postsynaptic currents in the ARC expressed *Kiss1.*
