## Supplementary figures and images for "Melanocortin-responsive Kiss1 neurons of the arcuate nucleus drive energy expenditure through glutamatergic signaling to the dorsomedial hypothalamus"

### Supplemental Figure 1

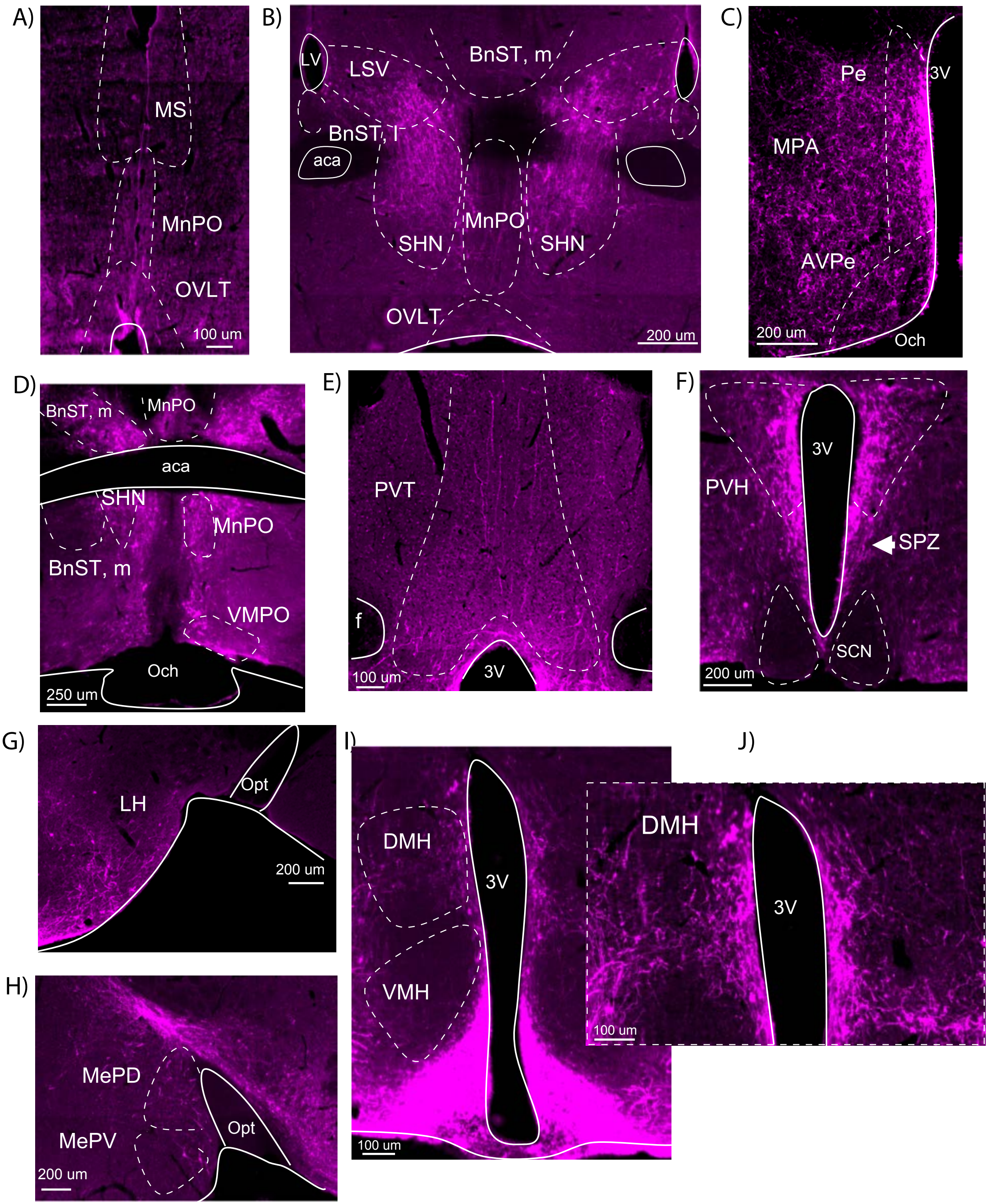

### Supplemental Figure 2

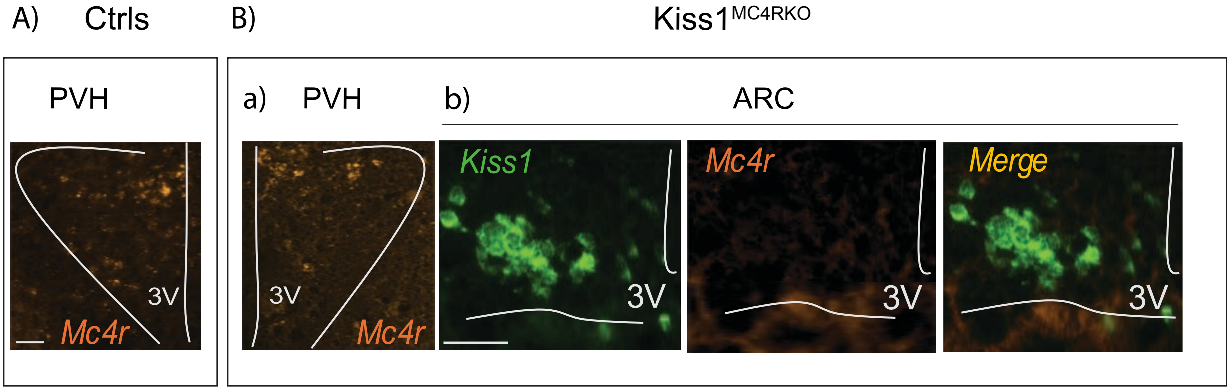

### Supplemental Figure 3

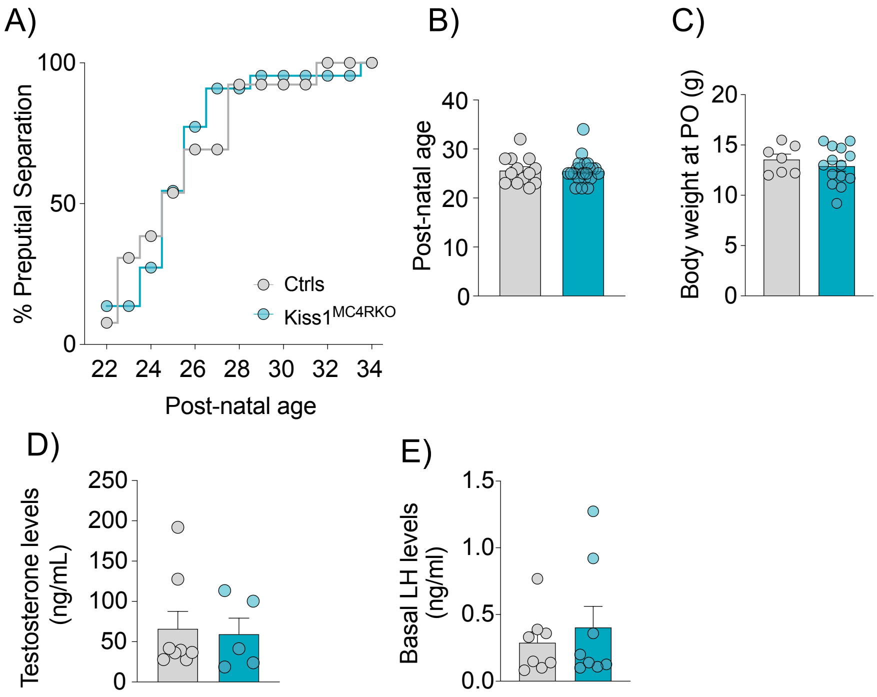

### Supplemental Figure 4

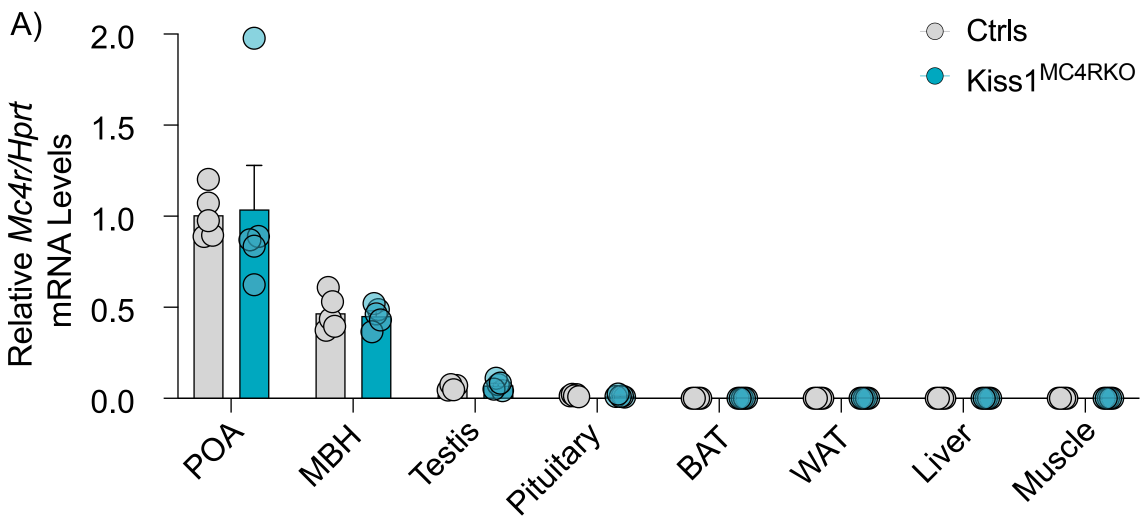

### Supplemental Figure 5

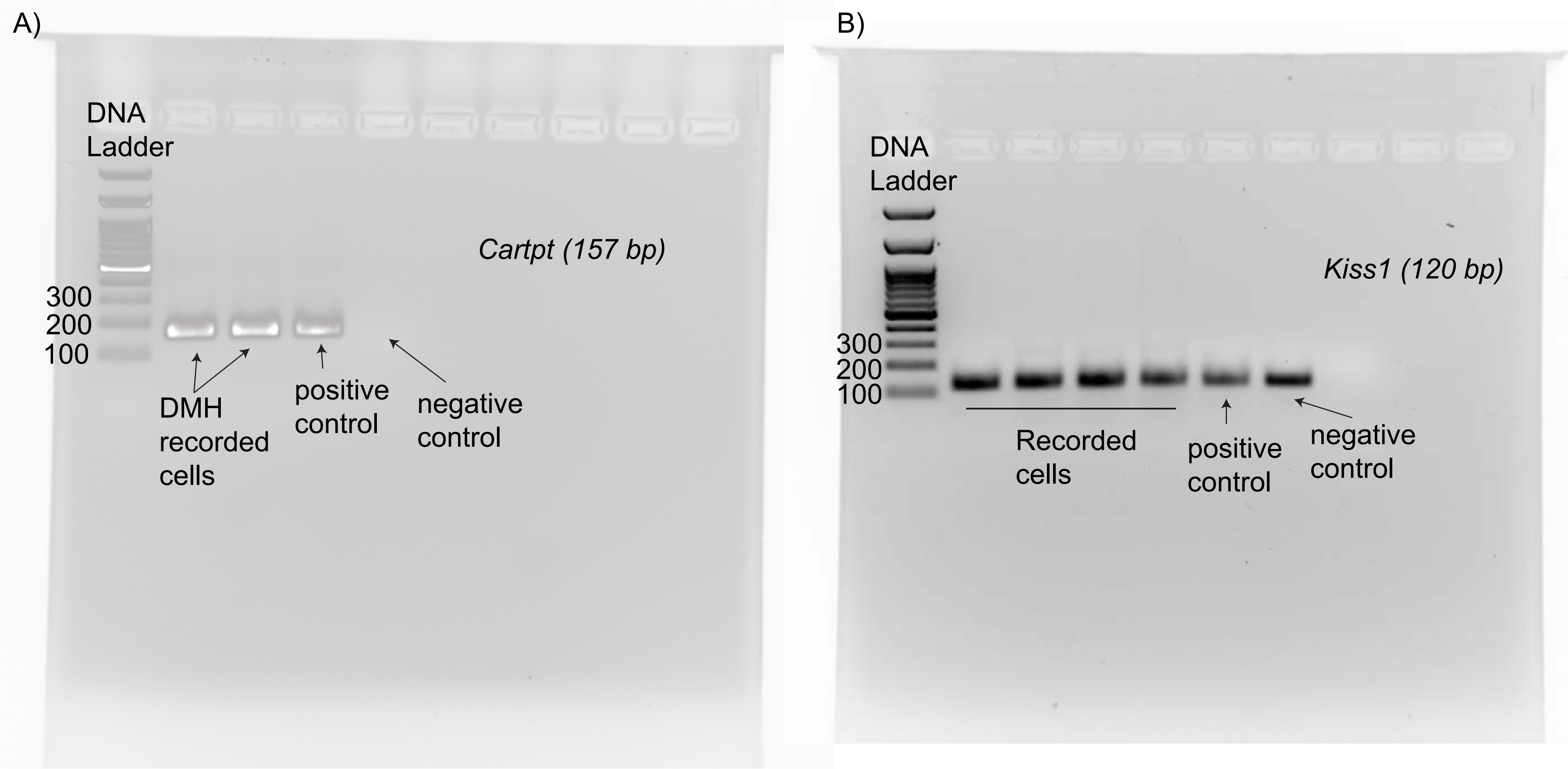
